# SINGLE CELL DISSECTION OF DEVELOPMENTAL ORIGINS AND TRANSCRIPTIONAL HETEROGENEITY IN B-CELL ACUTE LYMPHOBLASTIC LEUKEMIA

**DOI:** 10.1101/2023.12.04.569954

**Authors:** Ilaria Iacobucci, Andy G.X. Zeng, Qingsong Gao, Laura Garcia-Prat, Pradyumna Baviskar, Sayyam Shah, Alex Murison, Veronique Voisin, Michelle Chan-Seng-Yue, Cheng Cheng, Chunxu Qu, Colin Bailey, Matthew Lear, Matthew T. Witkowski, Xin Zhou, Airen Zaldivar Peraza, Karishma Gangwani, Anjali S. Advani, Selina M. Luger, Mark R. Litzow, Jacob M Rowe, Elisabeth M. Paietta, Wendy Stock, John E. Dick, Charles G Mullighan

## Abstract

Sequencing of bulk tumor populations has improved genetic classification and risk assessment of B-ALL, but does not directly examine intratumor heterogeneity or infer leukemia cellular origins. We profiled 89 B-ALL samples by single-cell RNA-seq (scRNA-seq) and compared them to a reference map of normal human B-cell development established using both functional and molecular assays. Intra-sample heterogeneity was driven by cell cycle, metabolism, differentiation, and inflammation transcriptional programs. By inference of B lineage developmental state composition, nearly all samples possessed a high abundance of pro-B cells, with variation between samples mainly driven by sub-populations. However, *ZNF384-*r and *DUX4-* r B-ALL showed composition enrichment of hematopoietic stem cells, *BCR::ABL1* and *KMT2A*-r ALL of Early Lymphoid progenitors, *MEF2D*-r and *TCF3::PBX1* of Pre-B cells. Enrichment of Early Lymphoid progenitors correlated with high-risk clinical features. Understanding variation in transcriptional programs and developmental states of B-ALL by scRNA-seq refines existing clinical and genomic classifications and improves prediction of treatment outcome.

## INTRODUCTION

Classification and risk stratification in B-cell acute lymphoblastic leukemia (B-ALL) primarily rely on integrating genomic and clinical factors^1–7^. Yet, diagnosis is defined based on the developmental stage of the disease revealed by the expression of immunophenotypic markers specific to a B-cell precursor stage. This distinction is crucial for differentiating B-ALL from other acute leukemias^1,8,9^. However, while the longstanding hypothesis for the origins of B-ALL implicates the Pro-B cell along a window of vulnerability during B cell receptor rearrangement, there is also evidence that origins may be heterogeneous^10–16^ and may sometimes arise from more primitive hematopoietic stem and progenitor cells (HSPC)^17^ or at more mature stages during immunoglobulin light chain rearrangement^18^. This heterogeneity challenges conventional diagnosis and outcome. Bulk transcriptome sequencing lacks the necessary resolution to examine distinct and heterogeneous subpopulations. By contrast, single-cell whole transcriptome (scRNA-seq) technology offers the potential to overcome this limitation, similar to its success in understanding acute myeloid leukemia (AML)^19,20^, potentially leading to a more precise and informative classification system for B-ALL.

Here, we performed scRNA-seq on 89 samples from 84 B-ALL patients to examine transcriptomic heterogeneity, developed a reference map of normal human B-cell development based on 130,085 cells from various sources, and found that single-cell transcriptome mapping in B-ALL identifies distinct developmental states associated with genomic alterations and clinical outcomes.

## RESULTS

### Single-cell RNA sequencing characterization of B-ALL

In this study we generated scRNA-seq data on 89 samples from 84 B-ALL patients, including different molecular subtypes (**Supplementary Tables 1-2**, **Fig. 1a** and **ED Fig. 1a**). Cells were annotated according to cell type into six broad hematopoietic cell types and in leukemic cells *versus* non-malignant cells, by applying the Inferred Copy Number Variation (inferCNV) algorithm (**Online Methods**) followed by manual cluster inspection^21^. We did not detect any CNVs in normal hematopoietic cells, while blast cells showed specific CNVs consistent with karyotype and whole genome sequencing (WGS) data (**Supplementary Table 1** and **ED Fig. 1b**). Moreover, single cell data was screened for driver sequence variations and gene fusions to confirm leukemic blast assignment especially in samples lacking chromosomal abnormalities. The gene expression values of cells from the same samples were aggregated in pseudobulk samples (N=89) and analyzed with 2,046 B-ALL bulk RNA-sequencing (RNA-seq) samples^22^ (**Supplementary Table 3**). This analysis demonstrated the robustness of scRNA-seq data to successfully cluster with their own corresponding leukemia subtype from bulk data (**Fig. 1b**). By Uniform Manifold Approximation and Projection (UMAP) analysis and visualization of scRNA-seq samples, normal cells from different samples and subtypes clustered together, while leukemic blasts formed distinct gene expression clusters (**Fig. 1c**).

**Figure 1.**
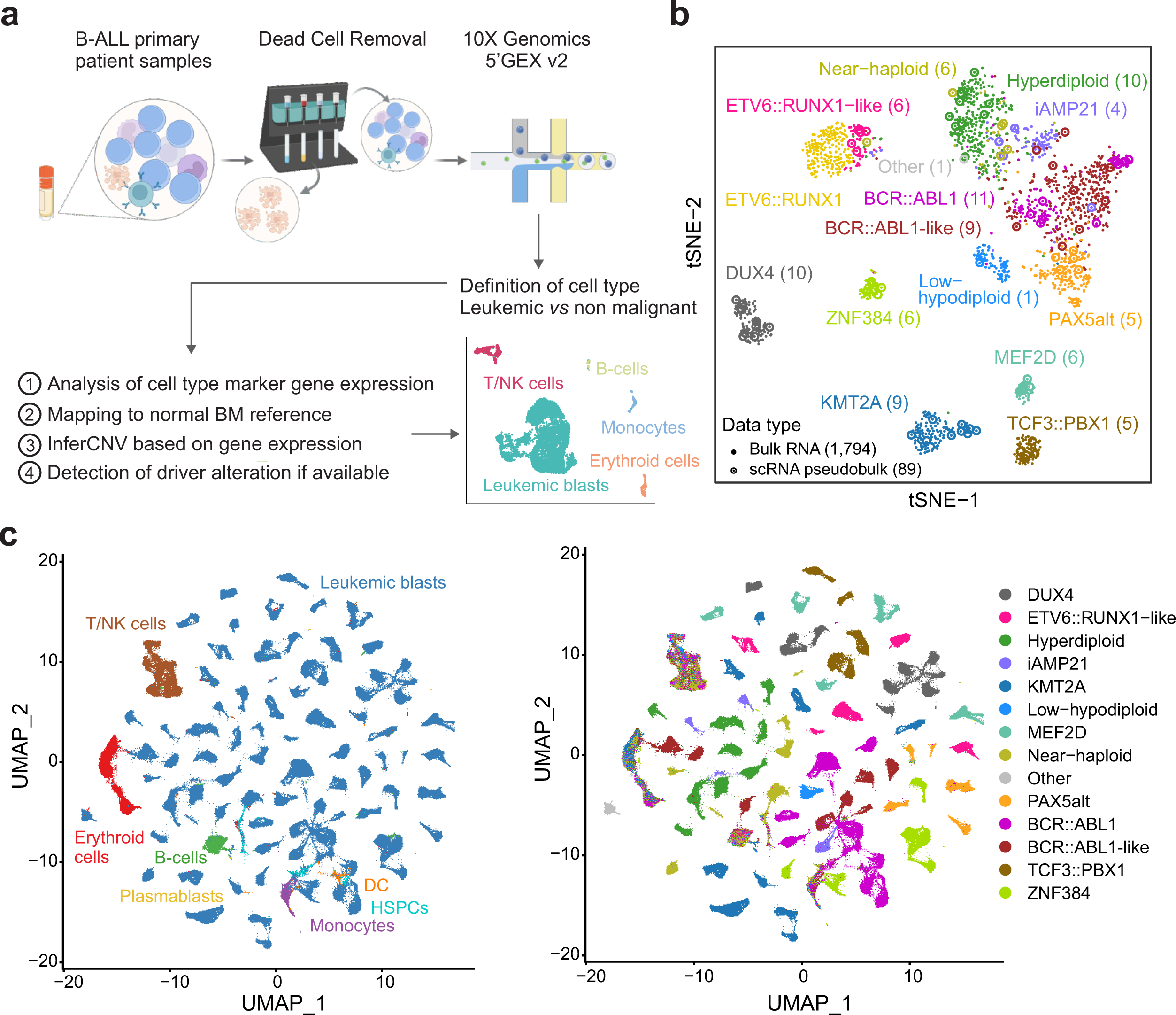
Single-cell RNA sequencing landscape of B-ALL. **a**) Schematic workflow for scRNA-seq of B-ALL samples. The figure was created by Biorender.com. **b**) t-distributed stochastic neighbor embedding (tSNE) plot showing gene expression profiling of 1,794 B-ALL bulk RNA-seq samples^22^ and 89 pseudobulk RNA-seq from this single-cell study. Each dot represents a sample and samples are colored by the leukemia subtype. In brackets are the number of pseudobulk RNA-seq samples. These samples clustered with bulk samples according to their specific subtype. For this analysis we only included subtypes that were represented in the scRNA-seq samples with the addition of *ETV6::RUNX1* samples to show co-clustering with *ETV6::RUNX1*- like. **c**) Uniform Manifold Approximation and Projection (UMAP) representation of cells (N=489,019) from the 89 single-cell RNA-seq data. Clusters of cells are colored by cell types (left panel) and leukemia subtype (right panel).

### Intra sample genetic and transcriptomic heterogeneity of malignant cells

To explore intra sample genetic heterogeneity, we used inferCNV . We found a relatively high complexity of CNV clonal patterns, with 43 (48.3%) samples exhibiting more than one CNV clone. Out of these, 16 samples had a minor clone comprising less than 3% of the cells, which was below the resolution of bulk sequencing (**Fig. 2a** and **Supplementary Table 4**). Notably, in hyperdiploid and near haploid B-ALL, chromosomal gains or losses had the same pattern almost universally in all leukemic cells from the same sample, consistent with early acquisition of aneuploidy in leukemogenesis^23^ (**Fig. 2b** and **ED Fig. 2a**). However, a minority of samples showed sequential evolution of aneuploidy: near-haploid SJHYPO117 showed progressive losses of chromosomes 8, 14, and 18, associated with distinct gene expression profiles, while two hyperdiploid samples harbored a minor clone with chromosomal gain in only 1% of the cells (chromosome 17 in SJBALL030276 and chromosome 12 in SJALL040100) ^23^ (**Figs. 2c-d** and **ED Figs. 2b-c**).

**Figure 2.**
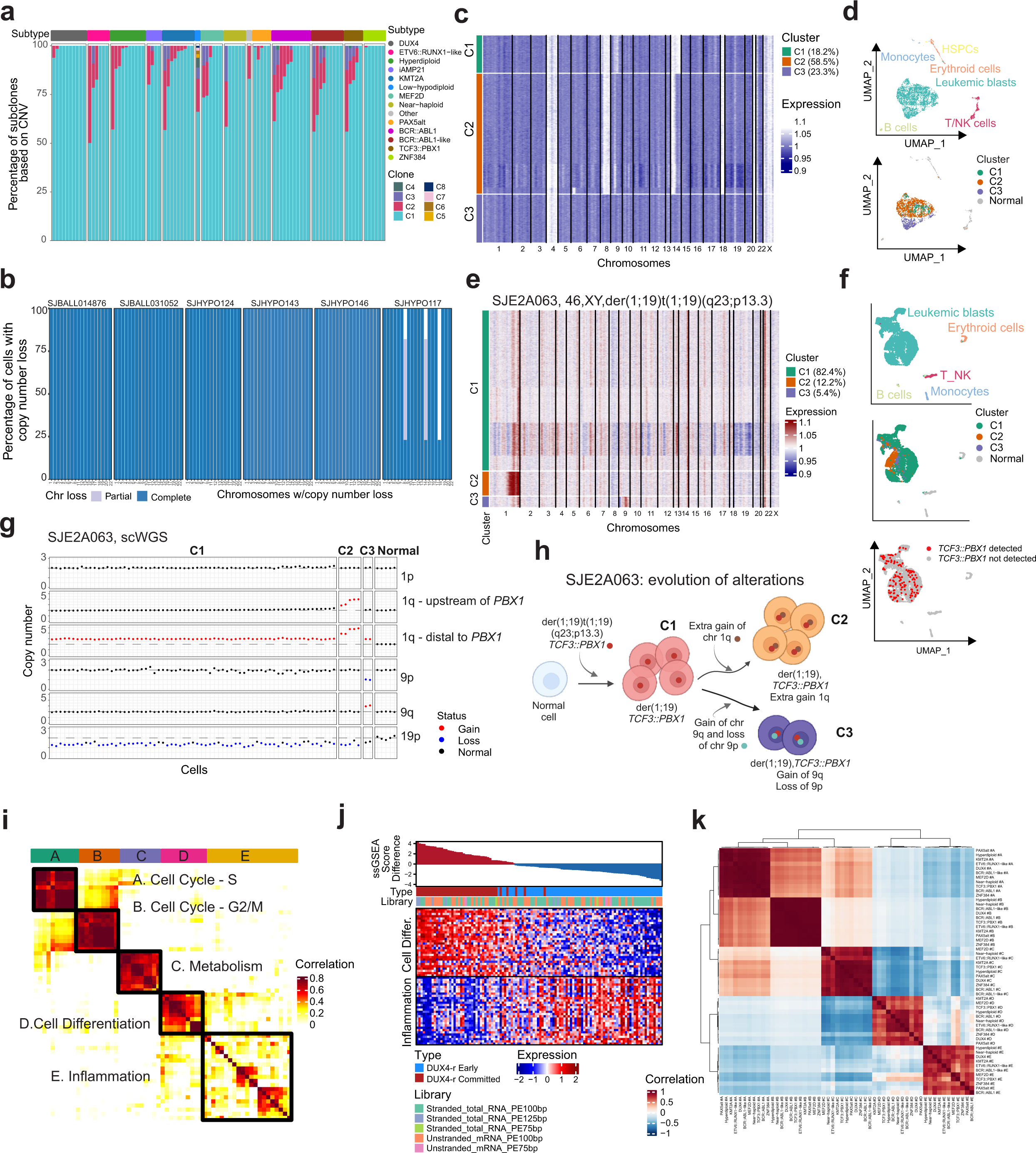
Single-cell genomic and transcriptomic heterogeneity. **a**) Bar plot showing copy number clonal heterogeneity in each sample and across subtypes. Number of clones ranged from 1 (N=46) to 8 (N=1, SJHYPO120). **b**) Bar plot showing partial or whole chromosome losses in near haploid B-ALL samples (N=6). **c**) InferCNV heatmap demonstrating total or partial chromosome losses in SJHYPO117. All leukemic cells showed loss of chromosomes 1, 2, 3, 5, 6, 7, 9, 10, 11, 12, 13, 15, 16, 17, 19, 20 and 22, suggesting that these alterations were earlier events arising in the founder clone. However, three clones differentiated for copy number changes of chromosomes 8, 14 and 18 and for gene expression profiles. While clone 1 and 2 clustered together, clone 3 with extra chromosomal losses showed a distinct cluster in the UMAP visualization. **d**) UMAP representation of cells from SJHYPO117. Clusters of cells are colored by cell type (upper panel) and inferCNV group (lower panel). Cells from cluster 3 without losses of chromosomes 8, 14p and 18 formed a distinct cluster (cluster 3). **e**) InferCNV heatmap from SJE2A063 (*TCF3::PBX1*) showing four CNV clusters and progressive gain of chromosome 1q. **f**) UMAP representation of cells from SJE2A063. Clusters of cells are colored by cell type (upper panel), inferCNV group (middle panel) and detection of *TCF3::PBX1* fusion (lower panel). **g**) Scatterplot showing copy numbers of chromosome (chr) 1p, chr 1q proximal (from centromere to *PBX1*; chr1: 144007038-164657928), chr 1q distal (from centromere to *PBX1*; chr1: 164657929-249250621), chr 9p (chr9:1-39500000), chr 9q (chr9:71000001-141213431) and chr 19p (chr19:1-1617900). Data are from scWGS in SJE2A063 performed on flow sorted blast (hCD45dim, CD19+) and normal (hCD45bright and CD3+) cells. Single cells were assigned to different clusters based on their copy number patterns. Dotted line indicates diploid DNA content. **h**) Cartoon created with Biorender.com and depicting the model of progressive gain of chromosome 1q and aberrations in chromosome 9 in SJE2A063 which occur in mutual exclusive clones. **i**) Heatmap showing the pairwise correlations of the sample level meta-programs derived from 10 *DUX4* patients using NMF. Clustering identified five coherent gene expression signatures of leukemic cells, including A - Cell Cycle (S), B - Cell Cycle (G2/M), C - Metabolism, D - Differentiation, and E - Inflammation. **j**) Heatmap showing gene expression pattern of the top signature 30 genes for subtype level programs 1 - Differentiation and 2 – Inflammation in 112 bulk RNA-seq B-ALL samples *DUX4*-r, which were ordered based on their enrichment score differences using ssGSEA. The sample information and library protocol were also annotated. **k**) Pairwise correlation of expression scores of transcriptional metaprograms from NMF defined separately in each subtype and applied across cells from all samples showing that cell cycle and metabolism expression programs are common across subtypes while those for cell differentiation and inflammation are subtype-specific.

Intratumoral variegation was also associated with chromosomal rearrangements encoding subtype-defining fusion oncoproteins, such as TCF3::PBX1, encoded by the t(1;19)(q23;p13) translocation. This rearrangement is commonly unbalanced, with duplication of 1q distal to *PBX1,* yet the mechanisms and patterns of associated copy number changes remain poorly understood and cannot be resolved using conventional methods. Cytogenetic analysis identified a der(19) in four of five *TCF3::PBX1* samples (**Supplementary Table 1**). However, integrated scRNA-seq and single cell whole genome sequencing (scWGS) revealed a more complex mechanism characterized not only by a gradual gain of chromosome 1 from *PBX1* to the telomere but also by the amplification of the entire chromosome 1q, encompassing *PBX1* (**Figs. 2e-h**, **ED Figs. 2d-e, Supplementary Tables 5-6**). This observation aligns with the acquisition of amplification in a region on chromosome 1 that includes *PBX1*, likely occurring on the chromosome not involved in the translocation. Supporting this mechanism, we noted amplification of the same region in a sample (SJEA067) with a balanced t(1;19) translocation. Single cell sequencing further unveiled clonal compositions indiscernible through bulk sequencing, revealing minor clones harboring additional chromosomal aberrations (**ED Figs. 2f-i)**. These data supported a mechanism by which *TCF3::PBX1* is an early event often accompanied by der(19) and amplification of the telomeric region distal to *PBX1* with additional copy number changes occurring in subclones. Thus, integrated single cell expression and copy number analysis reveal previously unrecognized clonal heterogeneity in B-ALL and a dynamic genetic landscape characterized by progressive accumulation of genetic alterations.

To explore intra sample transcriptional heterogeneity of leukemic cells, we first performed non-negative matrix factorization (NMF)^24^ of single cell gene expression data. Hierarchical clustering was performed on NMF programs from samples within the same subtype. This identified five consensus transcriptional meta programs in all subtypes independently of the genetic driver: cell cycle (S), cell cycle (G2/M), metabolism, cell differentiation, and inflammation (**Fig. 2i**, **ED Figs. 3, 4a** and **Supplementary Table 7**). Interestingly, cell differentiation and inflammation programs clearly separated *DUX4*-r samples in two groups consistent with bulk RNA-seq data^23^. According to cell differentiation program, we identified one class (“early”) with expression of early lymphocyte progenitor and myeloid genes (eg. *FLT3*, *CEBPA*) and inflammatory genes (eg. *TGFB*) and another class (“committed”) with expression of genes expressed in a more committed state of B-cell differentiation, such as *EBF1*, *CD79B*, *MME* (*CD10*) and *RAG1* (**ED Figs. 4b-e**). Flow cytometric analysis confirmed high expression of CD44 in *DUX4*-r “early” and CD10 in *DUX4*-r “committed” (**ED Figs. 4f-g**), providing diagnostic markers to prospectively define these two classes. Moreover, the defined cell differentiation and inflammation enrichment scores from scRNA-seq enabled the annotation of *DUX4*-r as either “early” or “committed” in an extended cohort of 112 *DUX4*-r B-ALL bulk RNA-seq (**Fig. 2j**), showing the power of these signatures to retrospectively assign class annotation to previously sequenced samples.

**Figure 3.**
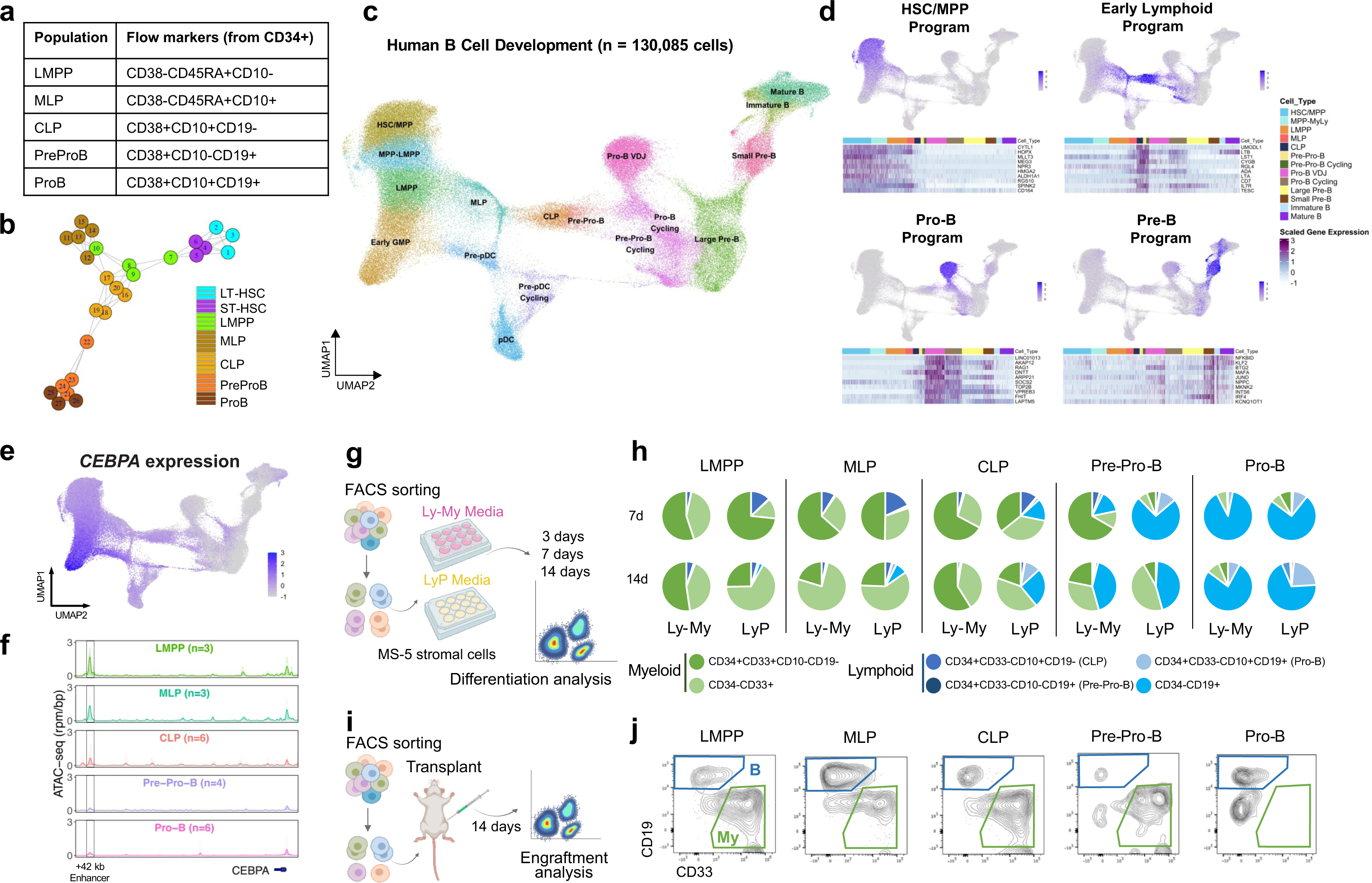
Molecular and functional characterization of human B cell development. **a**) FACS sorted cell populations along early human B cell development. **b**) Clustering of bulk RNA-seq profiles from sorted cell populations spanning B cell development in umbilical cord blood. Samples were clustered based on the expression of known human transcription factors (TFs). **c**) A single cell atlas of human B cell development, comprised of 130,085 single cell transcriptomes integrated from eight studies spanning fetal bone marrow, fetal liver, umbilical cord blood, pediatric bone marrow, and adult bone marrow tissues. Cell clustering and annotations were guided by bulk RNA-seq reference profiles from (**b**). **d**) cNMF signature analysis in normal B-lymphoid development. Signature strength of cNMF programs corresponding to HSC/MPP, Early Lymphoid, Pro-B, and Pre-B stages of B cell development are shown, together with an expression heatmap of 10 genes with the highest weights within each signature along B cell development. **e**) CEBPA regulon activity inferred through pySCENIC persists into the CLP stage of B cell development. **f**) Chromatin accessibility of an enhancer element +42kb from the CEBPA locus from sorted populations. **g-h**) *in vitro* differentiation assays of human lymphoid-primed multipotent progenitor (LMPP), multilineage progenitor (MLP), CLP, Pre-Pro-B, and Pro-B cells in myelo-lymphoid (My-Ly) or lymphoid (Ly) media. Two hundred cells from indicated cell populations were sorted and cultured on MS5 stroma cells for 3 (**ED Fig. 5**), 7 or 14 days in Ly-My or LyP medias (**Online Methods**) and analysed by flow cytometry. The proportion of each population is represented in pie charts. Cartoon created with Biorender.com. **i-j**) *in vivo* differentiation assays of human LMPP, MLP, CLP, Pre-Pro-B, and Pro-B cells. NSG mice were subject to intra-femoral injection of sorted populations and engraftment levels were analysed by flow cytometry two weeks later. Three independent experiments with a total n of 3-24 mice. Number of cells injected for each population is indicated in **ED Fig. 5**. Representative FACS plots and engraftment results are shown in **ED Fig. 6**.

**Figure 4.**
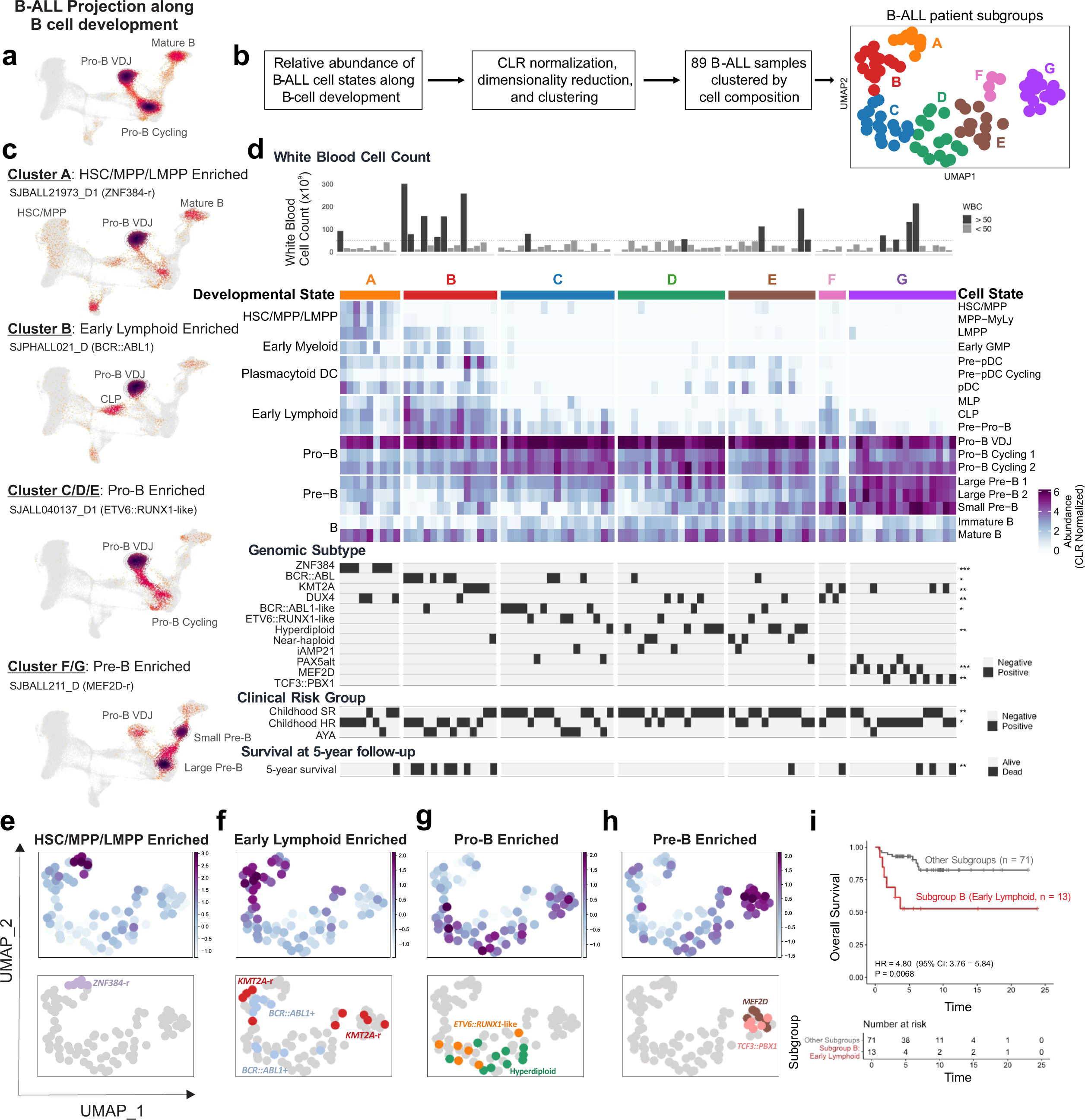
Heterogeneity in B cell developmental states implicated within B-ALL. **a)** Schematic outlining composition analysis of B-ALL samples along B cell development. B-ALL cells were projected onto the single cell atlas of human B-cell development and cell states were assigned based on the most transcriptionally similar healthy population. Samples were clustered based on the relative abundance of each cell state along B cell development. **b)** UMAP of 89 B-ALL samples grouped into six patient clusters based on cell composition. **c)** Projection results of B-ALL cells from representative patients within cell state composition-based clusters. **d)** Cell state composition-based clustering of 89 B-ALL samples. Each column represents a single sample, grouped into six clusters based on cell composition. Heatmap depicts normalized abundance of each mapped cell population for each sample, and annotations pertaining to white blood cell count, genomic subtype, clinical risk group, and 5-year survival are also provided for each sample. Statistical significance was determined by chi-squared tests between each annotation and the composition-based clusters: * < 0.05; ** < 0.01; *** < 0.001. **e**-**h**) UMAP clustering based on cell state composition from 89 B-ALL samples depicting abundance of broad developmental states spanning **e)** HSC/MPP/LMPP, **f)** Early Lymphoid, **g)** Pro-B, and **h)** Pre-B. Key genomic subtypes enriched for each developmental state are also depicted with their position along the UMAP embedding. **i**) Overall survival outcomes of patients in subgroup B compared to patients from other subgroups among samples collected from patients at initial diagnosis (n = 84). *P* values from likelihood ratio test are shown.

To explore transcriptional heterogeneity across different leukemia subtypes, we next analyzed how the expression scores of genes contributing to each metaprograms varied across different subtypes by pairwise correlation. This analysis revealed that transcriptional programs related to cell growth (S phase), cell division (G2/M phase), and metabolism were very similar across all subtypes (average r=0.97, r=0.99 and r=0.94, respectively) despite the differences in their genomic drivers. In contrast, cell differentiation and inflammation programs showed intermediate correlation (average r=0.65 and r=0.67, respectively), consistent with the observation that genes contributing to these two programs are different across different subtypes (**ED Fig. 4h**). Overall, this analysis revealed that cell differentiation composition and inflammation programs account for the transcriptomic heterogeneity across different leukemia samples.

### Characterization of human B cell development

Given the variation in cell differentiation composition from the NMF analysis, we sought to examine the ontogeny of B-ALL data through the lens of normal human B cell development. As B cell development has been extensively studied in mice^25^ and equivalent human data is lacking, we first created a refined map of human B cell development. We performed stringent sorting of immunophenotypically-defined cell populations of human B cell developmental stages from umbilical cord blood (CB; **Fig. 3a** and **ED Figs. 5a-b**). Clustering of sorted fractions from early B cell development confirmed their order along B cell differentiation **(Fig. 3b).**

**Figure 5.**
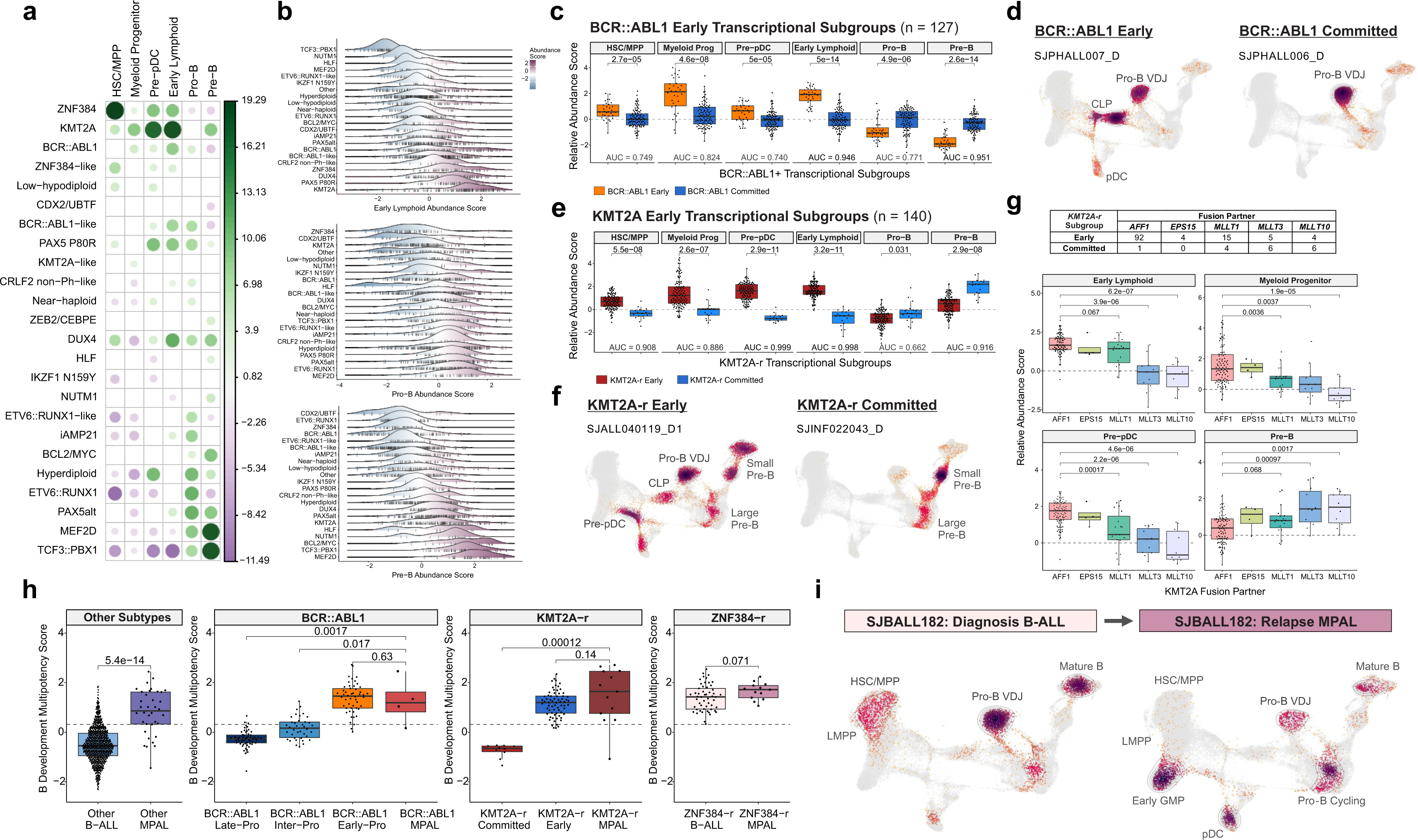
B-ALL developmental states refine existing genomic subgroups. **a)** Association between genomic subgroup and inferred abundance of B-ALL developmental states from 2,046 B-ALL patients profiled by RNA-seq. The magnitude of each association, quantified as the –log10 (p value), is depicted through the size and color intensity of each dot. The direction of the association is depicted through the color, wherein higher abundance is green and lower abundance is purple. Only associations with an FDR corrected *P*<0.05 are depicted. **b**) Ridgeplot comparing the inferred abundance of Early Lymphoid, Pro-B, and Pre-B states across genomic subtypes in B-ALL. **c**) Developmental state abundance explains transcriptional subtypes of *BCR::ABL1* B-ALL. **d**) Projection results from representative samples from the Early and Committed subtypes are shown. **e**) Developmental state abundance explains transcriptional subtypes of *KMT2A-r* B-ALL. **f**) Projection results from representative patients from the Early and Committed subtypes are shown. **g**) Association between *KMT2A* fusion partner and developmental state abundance. *KMT2A* fusion partners harbored by patients within each *KMT2A* subtype are shown alongside the inferred abundance of Early Lymphoid, Myeloid Progenitor, Pre-pDC, and Pre-B states.

We next utilized this resource to develop and annotate a single cell reference map of human B cell development onto which B-ALL blasts could be projected. We compiled scRNA-seq data from 8 normal cell datasets spanning fetal liver and bone marrow, cord blood, pediatric and adult bone marrow, focusing on cell states along B cell development and branch points to proximal lineages. Collectively, 130,085 cells were integrated across 90 donors, 5 tissues, and 3 technologies (**Supplementary Table 9** and **Fig. 3c**)^26–31^. Focused clustering and annotation guided by our reference transcriptomes from our sorted cell fractions allowed us to identify 17 cell states spanning human B cell development (**Fig. 3c**) while projection of surface protein levels from AbSeq data^32^ enabled immunophenotypic validation of our cell populations (**ED Figs. 5c-g and Supplementary Information)**.

To identify gene expression programs that underlie human B cell development, we applied NMF to our single cell map, identifying two cell cycle programs (S-phase and G2/M-phase) and seven stage-specific programs along B cell development (**Fig. 3d**, **ED Figs. 5h-i, Supplementary Table 10**). This included an “Early Lymphoid” program enriched in multi-lymphoid progenitors (MLPs) and common lymphoid progenitors (CLPs) which was characterized by high expression of *IL7R* and CD7 that encode known surface markers of common lymphoid progenitors in mice and humans (**ED Figs. 5j-k**)^33,34^. Flow cytometry analysis validated the stage-specific expression of surface IL7R and CD7 protein levels within MLPs and CLPs (**ED Figs. 5l-m**).

We next evaluated transcription factor (TF) activity, captured by enrichment of their target regulons, across lymphoid lineage commitment through SCENIC regulon analysis^35^. Unexpectedly, we found that the activity of the myeloid-associated TF CEBPA persisted into the CLP stage (**Fig. 3e**), despite previous characterizations of CLP as functionally restricted to B/NK lineages^36^. This was supported by bulk ATAC-seq on the sorted populations which revealed persisting accessibility of a CEBPA-regulating +42kb enhancer into the CLP stage (**Fig. 3f**). The retention of functional myeloid potential within CLPs but not Pro-B cells was confirmed through *in vitro* differentiation assays on MS5 stromal cells with either Lympho-Myeloid (Ly-My) or Lymphoid promoting (LyP) media (**Figs. 3g-h and ED Figs. 6a-c)** as well as *in vivo* xenotransplantation into NSG mice (**Figs. 3i-j** and **ED Figs. 6d-g)**. In further support of these findings, myeloid surface marker CD33 levels were higher in the CLP fraction compared to downstream B progenitor populations (**ED Fig. 6h**) and *in vitro* differentiation assays of CD33 positive and negative subsets of Early Lymphoid progenitors confirmed this association between CD33 and functional myeloid potential within the CLP fraction (**ED Figs. 6i-l**).

**Figure 6.**
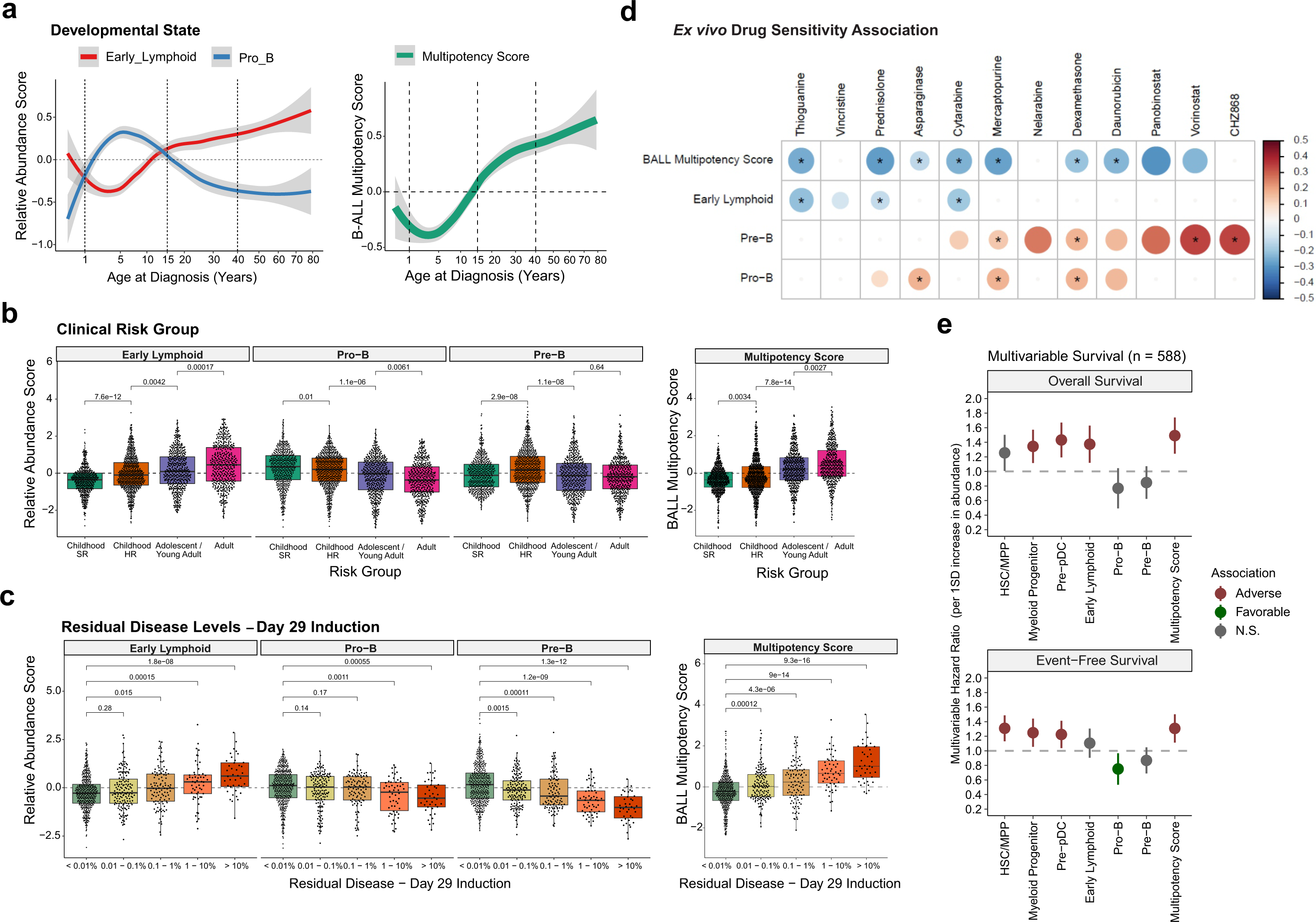
Clinical associations with B-ALL developmental states. **a)** Inferred abundance of Early Lymphoid and Pro-B states (left panel) and of Multipotency score (right panel) with age at B-ALL diagnosis. **b**) Association between inferred abundance of Early Lymphoid and Pro-B states (left panel) and of Multipotency score (right panel) with B-ALL clinical risk groups. **c**) Association between inferred abundance Early Lymphoid, Pro-B, and Pre-B states at B-ALL diagnosis with residual disease levels at day 29 of induction chemotherapy administration for 792 pediatric B-ALL patients with available information on residual disease levels. **d**) Association between Early Lymphoid, Pro-B, and Pre-B states with *ex vivo* drug sensitivity from 595 B-ALL samples profiled by bulk RNA-seq and *ex vivo* drug sensitivity screening for 18 therapeutic agents from Lee *et al* 2023 (ref^42^). Only agents with a significant p value < 0.05 are shown and those with the “*” symbol have FDR < 0.05. **e**) **d**) Association of B-ALL developmental states and of Multipotency score with survival outcomes within 588 pediatric B-ALL patients. Hazard ratios from multivariable Cox regression, adjusting for clinical risk group and genomic subtype, with overall survival and event-free survival are reported for each standard deviation increase in inferred abundance.

Altogether, we have developed a comprehensive atlas of cell states spanning human B cell differentiation from pre-natal and post-natal tissues. Furthermore, integration of molecular profiles with functional data from purified populations has revealed retention of myeloid lineage potential further into B-lymphoid development than previously expected, which may explain lymphoid-to-myeloid lineage switching in some B-ALL samples at disease relapse.

### Developmental states of B-ALL at single cell resolution

This map of human B cell development enabled precise mapping of B-ALL cells to developmental states, and dissection of heterogeneity, across the continuum of human B cell development.

Projection of B-ALL cells onto this normal reference revealed precise stages of human B cell development implicated in each patient’s disease (**Fig. 4a**). The “Pro-B VDJ” state, wherein heavy chain rearrangement occurs, was most frequently observed, and comprised at least 20% of a patient’s leukemic cells in 76 of 89 samples (**Supplementary Table 11**). By contrast, tumor samples varied in representation of other populations along B cell development. To simplify the many precise cellular states, we used NMF to condense co-occurring cellular states into broader categories, hereafter referred to as “developmental states” (**ED Figs. 7b-c** and **Supplementary Table 12**). We confirmed that B-ALL cells belonging to each developmental state retained stage-specific expression of cell identity programs from normal B cell development (**Supplementary Table 11** and **ED Figs. 8a-b**).

Clustering samples by cellular composition revealed seven patient subgroups (subgroups A-G; **Fig. 4a-d** and **ED Fig. 7a**) differing in enrichment of various B cell developmental states. Each subgroup was associated with distinct underlying genomic characteristics and clinical outcomes (**Figs. 4c-d** and **ED Figs. 7d-e**). A subgroup defined by high HSC/MPP abundance (subgroup A) was enriched for samples with rearrangements in *ZNF384* (**Fig. 4e**) that encode fusion oncoproteins enriched in lineage-ambiguous leukemia^37,38^, and a subset of *DUX4*-r ALL. Subgroup B was highly enriched with Early Lymphoid populations (MLP, CLP, Pre-Pro-B) and comprised of a subset of *BCR::ABL1*+ cases and a subset of *KMT2A*-r cases (**Fig. 4f**). A majority of samples including hyperdiploid, and *ETV6::RUNX1*-like genomic subtypes, was highly enriched with Pro-B populations (subgroups C, D, E; **Fig. 4g**). Last, a subset of samples (subgroups F, G) with high abundance of Pre-B cells, wherein light chain rearrangement occurs, was enriched for *TCF3::PBX1* and *MEF2D-*r subtypes (**Fig. 4h**).

Importantly, the early lymphoid-enriched subgroup B had inferior overall survival compared to other subgroups (HR = 4.80, *P*=0.0068) (**Fig. 4i**) and marginally worse event-free survival outcomes (HR = 2.67, *P* = 0.062) (**ED Fig. 7f**). Notably, this difference in overall survival between subgroup B and other subgroups persisted within *BCR::ABL1*+ patients (n = 10, *P* = 0.0074) but not within *KMT2A*-r patients (n = 8, *P* = 0.92) (**ED Fig. 7g**). No difference in event-free survival was observed between subgroup B and others within *BCR::ABL1* or *KMT2A*-r patients (**ED Fig. 7h**). Thus, projection of B-ALL cells across human B cell development reveals a new dimension of heterogeneity in B-ALL that complements but also refines genomic stratification in B-ALL.

### Associations between developmental state and genomic alterations in B-ALL

We next examined developmental state in a cohort of 2,046 transcriptomes from childhood and adult ALL. We used our normal and malignant scRNA-seq data to infer B-ALL developmental state in the bulk RNA-seq cohort using 145 marker genes (**ED Figs. 8a-f**, **Supplementary Tables** 3 and **13-15**). This confirmed associations between developmental state and genomic subtype identified from scRNA-seq composition analysis, including HSC/MPP enrichment in *ZNF384*-r; early lymphoid enrichment in subsets of *KMT2A-*r, *BCR::ABL1*, and *BCR::ABL1*-like; Pre-B enrichment in *MEF2D*-r and *TCF3::PBX1*; and Pro-B enrichment with hyperdiploid and *ETV6::RUNX1-*like ALL (**Figs. 5a-b** and **ED Figs. 8g-i**). We identified associations between developmental state abundance with specific gene fusions and specific driver mutations. Notably, mutations involving *FLT3, NRAS*, and *IKZF1* were associated with increased abundance of earlier developmental states while mutations involving *RAG1/2,* and *CDKN2A/B* were associated with an increased abundance of more committed states (**ED Fig. 8k**).

We next asked whether developmental state information could be used to refine existing genomic subtypes. Our NMF analysis demonstrated that transcriptomic subtypes within *BCR::ABL1+* B-ALL^10^, *DUX4*-r B-ALL, and *KMT2A*-r B-ALL^23^ captured distinct cell differentiation states (**Figs. 2j-k**). Indeed, developmental state abundance confirmed differences between Early and Committed transcriptional subtypes of *DUX4*-r B-ALL, wherein the *DUX4-r* Early subgroup was highly enriched for HSC/MPP (AUC = 0.975) and Early Lymphoid (AUC = 0.850) abundance (**ED Figs. 9a-b**). Developmental state abundance also explained differences between previously reported Early-Pro, Inter-Pro, and Late-Pro transcriptional subgroups of 54 *BCR::ABL1* B-ALL samples^10^ and 127 *BCR::ABL1* B-ALL samples from our cohort (**ED Figs. 9c-g**). Simplifying these three classes into Early (Early_Pro) and Committed (Inter_Pro and Late_Pro) subgroups, Early Lymphoid abundance explained differences between transcriptional subgroups with high concordance (our data: AUC=0.946; Kim et al^10^: AUC=0.953) (**Fig. 5c** and **ED Figs. 9f-g)**. We confirmed these differences in cell composition between samples in each subgroup when projecting single cell samples on the human B cell map (**Fig. 5d**).

Last, we asked whether developmental state subtyping could increase our understanding of *KMT2A*-r B-ALL. In B-ALL, *KMT2A*-r has recently been shown to involve ELP-like states with stemness properties^39,40^. Transcriptional subgroups within 140 *KMT2A*-r patients can be classified as Early and Committed. The “Early subtype” was highly enriched for Early Lymphoid (AUC = 0.998), Pre-pDC (AUC = 0.999), and HSC/MPP (AUC = 0.908) cells. By contrast, the “Committed subtype” was highly enriched for Pre-B (AUC = 0.916) (**Fig. 5e**) cells. These differences were also evident from scRNA-seq profiles (**Fig. 5f**), suggesting heterogeneity in the transforming cell type underlying these two *KMT2A*-r subsets. Notably, these subgroups were associated with distinct *KMT2A* fusion partners (**Fig. 5g**), wherein *KMT2A::AFF1* fusions were associated with higher Early Lymphoid, Pre-pDC, and Myeloid Progenitor abundance while *KMT2A::MLLT3* and *KMT2A::MLLT10* fusions were associated with higher Pre-B abundance (**Fig. 5g**).

Given our observation that CLPs retain functional myeloid potential (**Fig. 3g-j**), and given that this Early Lymphoid developmental state, which incorporates MLP, CLP, and Pre-Pro-B, is highly enriched within the Early subgroup within each of the *BCR::ABL1*, *DUX4-r*, and *KMT2A-r* B-ALL subtypes, we asked whether the Early subgroup may express any myeloid signatures. Interestingly, each of these genomic subgroups with Early Lymphoid dominance showed overexpression of *CEBPA* compared to their Pro-B enriched counterparts (**ED Fig. 9h**). These data suggest that, in line with their counterparts from normal hematopoiesis, some B-ALL cells from patient samples belonging to these Early subgroups may retain latent myeloid potential. This suggests that the cell of origin of genomic subtypes characterized by lineage variability or ambiguity may be as late as a committed lymphoid progenitor that retains myeloid potential, rather than necessarily being an uncommitted hematopoietic stem cell or progenitor

### Early-subtype B-ALLs derive from multipotent progenitor cells

Given the observed enrichment of HSC/MPP and Early Lymphoid developmental states within the Early subgroups of B-ALL subtypes and the concomitant overexpression of myeloid regulator *CEBPA*, we hypothesized that these B-ALL subgroups may originate through leukemic transformation of a multipotent cell population, as previously demonstrated for *ZNF384-*r leukemia^37^. To examine this, we developed a B cell development Multipotency Score, which was positively associated with HSC/MPP, Early Lymphoid, Myeloid Progenitor, and Pre-pDC abundance and negatively associated with Pro-B and Pre-B abundance. We confirmed that this score was enriched across normal HSPCs compared to committed B-lymphoid precursors (**ED Fig. 9i)**. Within the malignant context, we compared bulk RNA-seq from B-ALL patients with those with B/Myeloid MPAL^37,41^ and confirmed that Multipotency Score levels in MPAL were significantly higher than in B-ALL (**Fig. 5h**).

However, these distinctions between B-ALL and MPAL were more nuanced in the context of *BCR::ABL1*+, *KMT2A*-r, and *ZNF384*-r genomic subtypes, which span both diseases. As expected, Multipotency Scores were significantly higher in Early subgroups compared to Committed subgroups within *BCR::ABL1*+ and *KMT2A*-r B-ALL subtypes (**Fig 5h**). However, Multipotency Scores within Early subgroups of *BCR::ABL1*+ and *KMT2A*-r B-ALL were similar to those of MPAL with the same genomic alterations (**Fig 5h**). Finally, *ZNF384*-r B-ALL, which exhibits the most HSC/MPP enrichment of all B-ALL subtypes, had comparable Multipotency Scores to *ZNF384*-r MPAL, highlighting their similar cellular origins.

This suggests that a subset of B-ALL samples may share a multipotent progenitor origin with MPAL samples containing the same genomic alterations, and suggests latent myeloid potential within *ZNF384*-r B-ALL and Early subgroups of *BCR::ABL1*+ and *KMT2A*-r B-ALL. Indeed, among B-ALL patients profiled by scRNA-seq we identified one diagnosis-relapse pair wherein a *ZNF384* B-ALL patient (SJBALL182) with high HSC/MPP involvement at diagnosis presented with both lymphoid and myeloid blasts at relapse. In the relapse sample for this patient, we observed the persistence of the primitive population alongside the expansion of downstream myeloid progenitors (**Fig. 5i**).

### Developmental state associations with leukemia outcome

Within the 2,046 B-ALL patient cohort, developmental state abundance was associated with age at diagnosis: leukemic samples from children had high leukemic cell Pro-B abundance, by contrast, those diagnosed in infancy or adulthood had higher abundance of more primitive cell states including Early Lymphoid and HSC/MPP cells (**Fig. 6a and ED Figs. 10a-d**). Furthermore, within childhood, high-risk disease was associated with high Early Lymphoid abundance and low Pro-B and Pre-B abundance compared to standard-risk disease, among other differences (**Fig. 6b and ED Figs. 10e-g**). Thus, higher risk disease and older age is associated with early multipotent stages of B cell development, likely reflecting age-dependent differences in susceptibility to leukemic transformation along human B cell development.

Among a subset of pediatric patients (N= 1,197) with measurable residual disease (MRD) information at day 29 of induction therapy, high Early Lymphoid abundance and low Pro-B and Pre-B abundance was observed in MRD positive patients compared to MRD negative patients **(ED Fig. 10h).** These populations were also associated with extent of residual disease among patients who were MRD positive (**Fig. 6c and ED Fig. 10i**). This suggests chemosensitivity of Pro-B and Pre-B cells and possible chemo-resistance of Early Lymphoid cells, which was confirmed through analysis with *ex vivo* drug sensitivity of 18 therapeutic agents from 595 B-ALL samples profiled by bulk RNA-seq^42^. This analysis revealed positive association of Pro-B and Pre-B abundance, and negative association of Early Lymphoid abundance, with *ex vivo* sensitivity to common chemotherapeutic agents used in B-ALL (**Fig. 6d**), in line with their associations to MRD levels.

Critically, developmental state was associated with survival outcomes. Multivariable analysis controlling for clinical risk group and genomic subtype within 588 pediatric B-ALL patients revealed that higher Early Lymphoid abundance was independently associated with adverse overall survival outcomes, while higher Pro-B abundance was independently associated with favorable event-free survival outcomes (**Fig. 6e** and **ED Fig. 10j**). These survival associations were more nuanced within adult B-ALL. Within a cohort of 214 genetically diverse B-ALL patients that did not include *BCR::ABL1+* B-ALL, no significant associations were identified between outcomes and developmental state abundance by multivariable analysis (**ED Figs. 9k-l**). However, within a cohort of adult *BCR::ABL1+* B-ALL patients (n = 41), higher Early Lymphoid abundance was associated with adverse overall and event-free survival outcomes (**ED Fig. 10m**).

## DISCUSSION

Here we provide multiple insights into the functional and cellular heterogeneity that exists in B-ALL by examining genetic complexity through the lens of intratumor transcriptional heterogeneity and B-cell development state. This work exceeds prior scRNA studies in B-ALL where only few samples and subtypes were analyzed^11,43^.

First, the analysis of genetic heterogeneity by inferCNV highlighted the temporal and quantitative patterns of CNV evolution in B-ALL and enabled the identification of minor clones with specific CNV patterns, which were not revealed by bulk sequencing data, for example the progressive chromosome 1q gain in *TCF3::PBX1* ALL. Moreover, it revealed the genetic evolutionary nature of aneuploidies in B-ALL with most samples often exhibiting the same chromosomal changes in all cells supporting a single event as origin of aneuploidy. However, it also revealed in a minority of cases a more complex clonal evolution pattern with alterations evolving progressively over time. Overall, these findings improve our understanding of the genetic intra tumor heterogeneity of B-ALL.

Second, this study provides a framework for understanding normal and malignant B-cell development by integrating functional analysis of meticulously sorted cell populations throughout human B-cell development with single-cell transcriptomics of B-ALL. A key finding from this analysis was the demonstration that CLPs retain myeloid potential, a finding that challenges our previous understanding of lymphoid lineage restriction and has important implications for lineage specification and promiscuity in B-ALL. Notably, we identified distinct developmental states within B-ALL, ranging from stem cells to more committed stages, which are associated with genomic drivers and provide a finer classification of B-ALL patients. For example, stem cells were enriched in *ZNF384*-r B-ALL, Early Lymphoid cells in a subset of *KMT2A*-r and *BCR::ABL1* B-ALL, while more differentiated lymphoid progenitors were enriched in *TCF3::PBX1* and *MEF2D* subtypes. Findings from our study also demonstrated that previously described subgroups of B-ALL, such as *KMT2A*-r, *DUX4*-r, and *BCR::ABL1* subtypes, can be explained by categorizing patients into Early/Multipotent and Committed subgroups based on developmental states. This sheds light on the shared cellular origins of these subgroups, which may explain the observed heterogeneity in the disease and its clinical outcomes.

Third, early multipotent stages of B cell development were associated with higher risk disease and older age at diagnosis, suggesting that the developmental context might influence the disease’s aggressiveness and response to treatment. Furthermore, association between these states and higher levels of MRD as well as inferior overall survival, independent of genomic subtype in pediatric patients, suggests inherent differences in sensitivity to conventional B-ALL chemotherapy between developmental states.

Critically, cell type composition may represent a risk factor to consider preventing lineage switch under specific-lineage targeted therapies and thus important to detect. For example, lineage switch from ALL to AML, especially under pressure of B-cell–specific immunotherapy, has been reported for patients with *KMT2A*-r and *BCR::ABL1*^44–47^ and been attributed to the existence of a primitive hematopoietic stem/progenitor cell as the cell of origin retaining myeloid potential^39^. Interestingly, all three subtypes with the “early” developmental state showed overexpression of *CEBPA*, whose activity we demonstrated by integrated genomic and functional studies to persist into the CLP stage. This finding may provide a biological explanation to the overexpression of *CEBPA* which has been previously reported in B-ALL subtypes, such as *ZNF384*-r, *DUX4*-r and PAX5 P80R, undergoing monocytic lineage switch with a gradual decrease in CD19 expression accompanied by a gradual increase in the expression of at least one monocytic marker^48^. This biological change in phenotype can result in discordant MRD levels determined by flow cytometry and molecular PCR-based assays and affects the availability of CD19 as a therapeutic target.

Finally, given the clinical implications of cell developmental states in B-ALL, especially with the emergence of new lymphoid-targeted therapies and the potential for phenotype shift as mechanism of escape, we developed a multipotency score, which was positively associated with multipotent cell populations and negatively associated with committed B-cell states. Notably, *ZNF384*-r B-ALL, characterized by stem cell enrichment, had a similar multipotency score to *ZNF384*-r MPAL, suggesting shared cellular origins and justifying the need for a classification based on genomic/cellular state features rather than immunophenotypic criteria alone.

In conclusion, elucidation of distinct developmental states in B-ALL improves our understanding of this leukemia, providing new insights into its molecular heterogeneity, cell origins, and impact on patient outcomes. Dissecting genetic subtypes into cell developmental subgroups may help guide treatment strategies to target specific cellular origins and developmental states, ultimately improving diagnosis and the effectiveness of B-ALL therapies.

## DATA AVAILABILITY

Single-cell RNA and DNA sequences have been deposited in the European Genome-phenome Archive (EGA) under accession number EGAS00001007512 and at Alex’s Lemonade Stand Foundation for Childhood Cancer single cell portal (https://scpca.alexslemonade.org/projects/SCPCP000008). Feature-barcode matrix, feature and barcode sequences from cellranger output of single-cell RNA-seq data have also been deposited to GEO under accession number GSE241405. RNA-seq from immunophenotypically sorted populations have been deposited to GSE125345.

## ACKNOWLEDGMENTS

This study was supported by the American Lebanese Syrian Associated Charities of St. Jude Children’s Research Hospital; the Alex’s Lemonade Stand Foundation for Childhood Cancer (C.G.M.); the National Institutes of Health, National Cancer Institute grants P30 CA021765 (St. Jude Children’s Research Hospital Comprehensive Cancer Center support grant) and R35 CA197695 (C.G.M.); a St. Baldrick’s Foundation Robert J. Arceci Innovation Award (C.G.M.); the Henry Schueler 41&9 Foundation (C.G.M.); University of Toronto MD/PhD studentship award (A.G.X.Z.); Princess Margaret Cancer Foundation (J.E.D.); Ontario Institute for Cancer Research through funding provided by the Government of Ontario (J.E.D.); Canadian Institutes for Health Research RN380110-409786 (J.E.D.); International Development Research Centre Ottawa Canada (J.E.D.); Canadian Cancer Society 703212 (J.E.D.); Terry Fox New Frontiers Program project grant 1106 (J.E.D.); University of Toronto’s Medicine by Design initiative with funding from the Canada First Research Excellence Fund (J.E.D.); The Ontario Ministry of Health (J.E.D.); Canada Research Chair (J.E.D.). Supported by SWOG NCTN grant: U10CA180888. Supported in part by the National Cancer Institute of the National Institutes of Health under award numbers: U10CA180820 and UG1CA189859. The content is solely the responsibility of the authors and does not necessarily represent the official views of the National Institutes of Health.

## AUTHOR CONTRIBUTIONS

I.I.: designed and led the study, performed experiments, analyzed data and wrote the manuscript; A.G.X.Z: contributed to study design, developed B-cell developmental maps, analyzed data and wrote the manuscript; Q. G.: analyzed data and wrote the manuscript; L. G.P.: performed functional experiments and wrote the manuscript; P.B.: contributed to perform single-cell experiments; S.S.: analyzed data; A.M.: analyzed data; V.V.: performed and analyzed data; M.C.S.: performed and analyzed data; C.C.: reviewed statistical analysis; C.Q.: performed data analysis; C.B.: contributed to sample collection; M.L.: provided clinical data; M.T.W.: contributed to data analysis; X.Z.: data visualization; A.Z.P.: data visualization; K.G.: data visualization; A.S.A.: provided clinical data; S.M.L.: provided clinical data; M.R.L.: provided clinical data; J.M.R.: provided clinical data; E.M.P.: provided clinical data; W.S.: provided clinical data; J.E.D.: contributed to study design, data analysis and revised the manuscript for final approval; C.G.M.: contributed to study design and revised the manuscript for final approval. All authors contributed to manuscript writing and gave final approval.

## EXTENDED DATA FIGURE LEGENDS

**Extended Data Figure 1.**
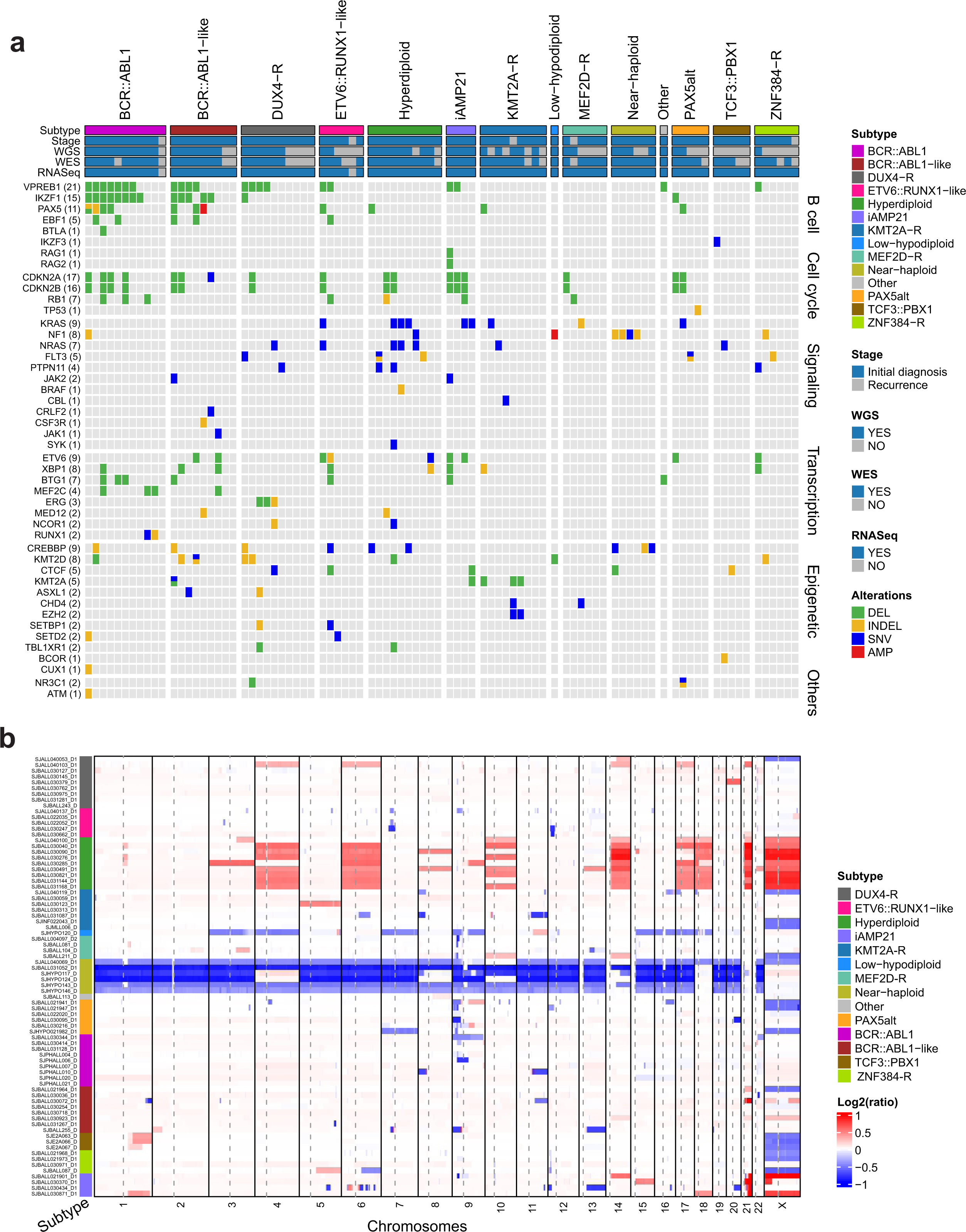
Molecular landscape of samples analyzed by scRNA-seq. **a**) For each sample single nucleotide variations (SNV), small indels (INDEL), focal deletions (DEL) and amplifications (AMP) are shown. Samples are grouped by their molecular subtype and for each sample, disease stage (initial diagnosis or recurrence) and analysis by WGS/WES/RNA-seq are shown. **b**) Copy number changes are depicted in red for copy number gains and in blue for copy number losses. Only samples with genomic data available are included.

**Extended Data Figure 2.**
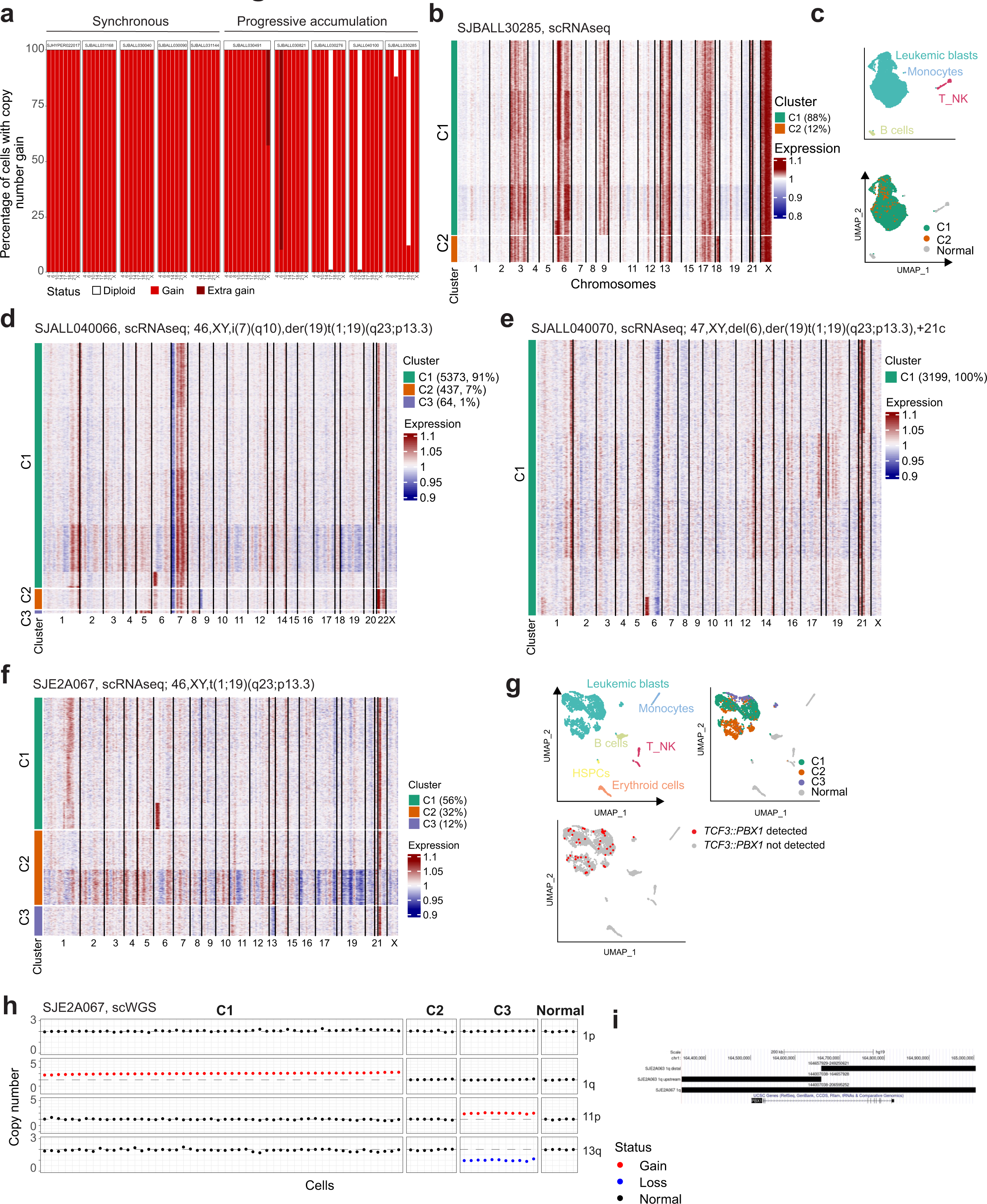
Genetic heterogeneity elucidated by inferCNV. **a**) Bar plot showing inferred copy number in samples with hyperdiploid. **b**) InferCNV heatmap demonstrating total or partial chromosome gains in SJBALL030285_D1. **c**) UMAP representation of cells from SJBALL030285_D1. Clusters of cells are colored by cell type (upper panel) and inferCNV group (lower panel). **d-e**) InferCNV heatmap from SJALL040066 (**d**) and SJALL040070 (**e**) samples with *TCF3::PBX1* fusion derived from the unbalanced translocation der(19)t(1;19)(q23;p13.3) as documented by cytogenetic analysis. The heatmap shows in SJALL040066 (**d**) three distinct CNV clusters sharing gain of 1q region from *PBX1* to telomere (chr1:164657929-249250621) and chr7p/7q gain but with acquisition of additional specific alterations which were undetectable by bulk sequencing approaches: loss of chr 9p and gain of chr 22 in clone 2 (C2); gain of chr 5 and chr 8 in clone 3 (C3). The heatmap shows in SJALL040070 (**e**) one CNV clone with gain of 1q region from *PBX1* to telomere (chr1:164657929-249250621). **f**) InferCNV heatmap from SJE2A067 with *TCF3::PBX1* fusion derived from balanced translocation t(1;19)(q23;p13.3) as documented by cytogenetic analysis. InferCNV analyses revealed 3 copy number clusters having distinct gene expression profiles (as shown in panel **g**): clone 2 (C2) is the founder clone; clone 1 (C1) is characterized by a gain of chr 1 upstream of *PBX1* (chr1: 144007038-164657928); clone 3 (C3) shows loss of chr 13. **g**) UMAP representation of cells from SJE2A067. Clusters of cells are colored by cell type (upper left panel), inferCNV group (upper right panel) and detection of *TCF3::PBX1* fusion (lower left panel). **h**) Scatterplot from scWGS in SJE2A067showing copy numbers of chr 1p, 1q regions upstream and downstream of *PBX1*, 11p, 13q and 19p. Dotted line indicates diploid DNA content. This analysis revealed multiple distinct clones not detectable by bulk sequencing demonstrating the complex patterns of genetic evolution in samples with *TCF3::PBX1* fusions. i) Screenshot from Genome Brower showing the locations of alterations on chr1 in SJE2A063 with amplification of regions both upstream and downstream *PBX1* as derived from two independent events and in SJE2A067 with a single alteration encompassing *PBX1* gene.

**Extended Data Figure 3.**
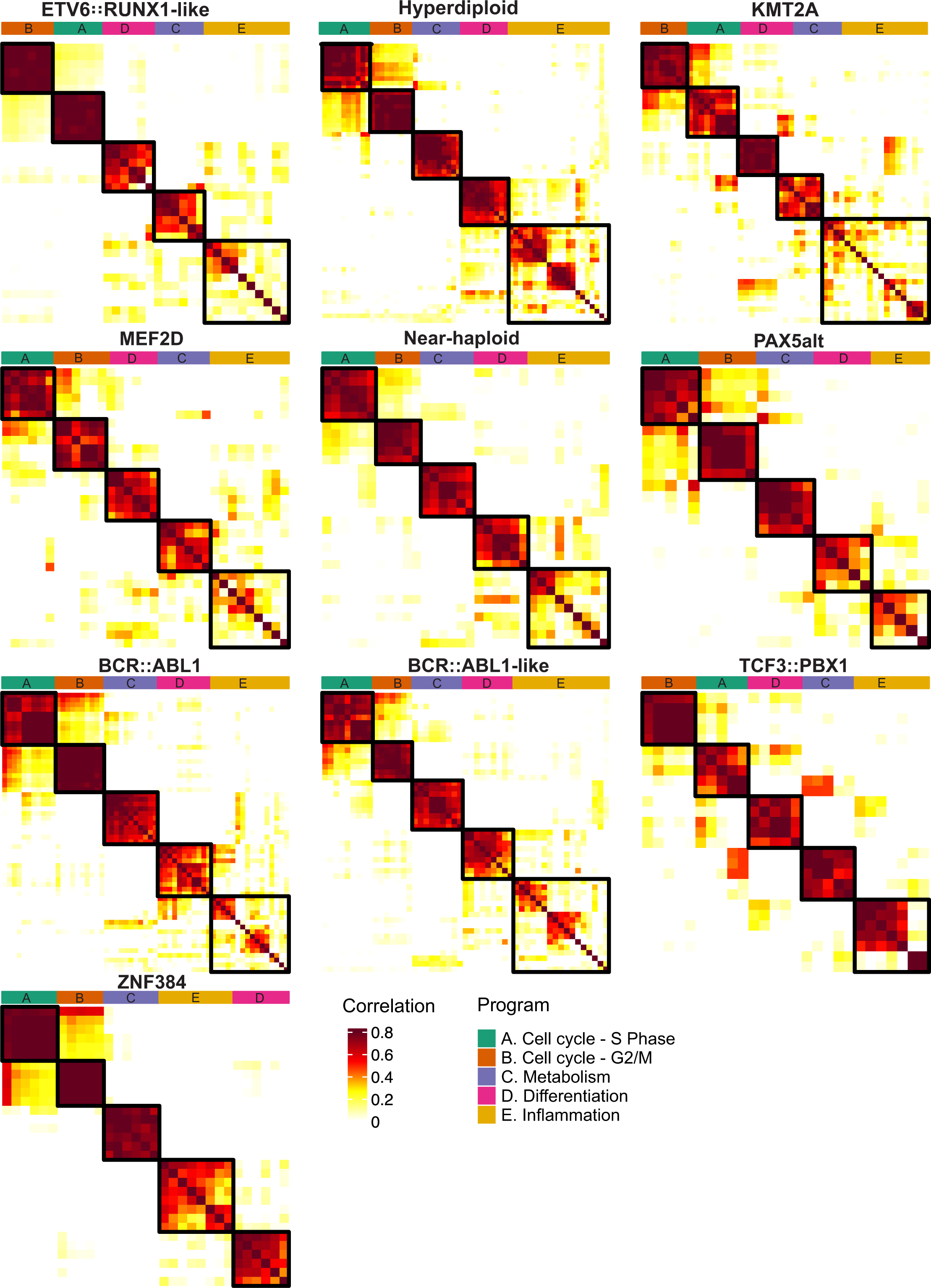
Transcriptional programs in B-ALL samples. Heatmap showing the pairwise correlations of the sample level meta-programs using consensus non-negative matrix factorization (cNMF). Each panel includes samples from the same subtype. Clustering identified five coherent malignant gene expression signatures, including A - Cell Cycle (S), B - Cell Cycle (G2/M), C - Metabolism, D - Differentiation, and E - Inflammation.

**Extended Data Figure 4.**
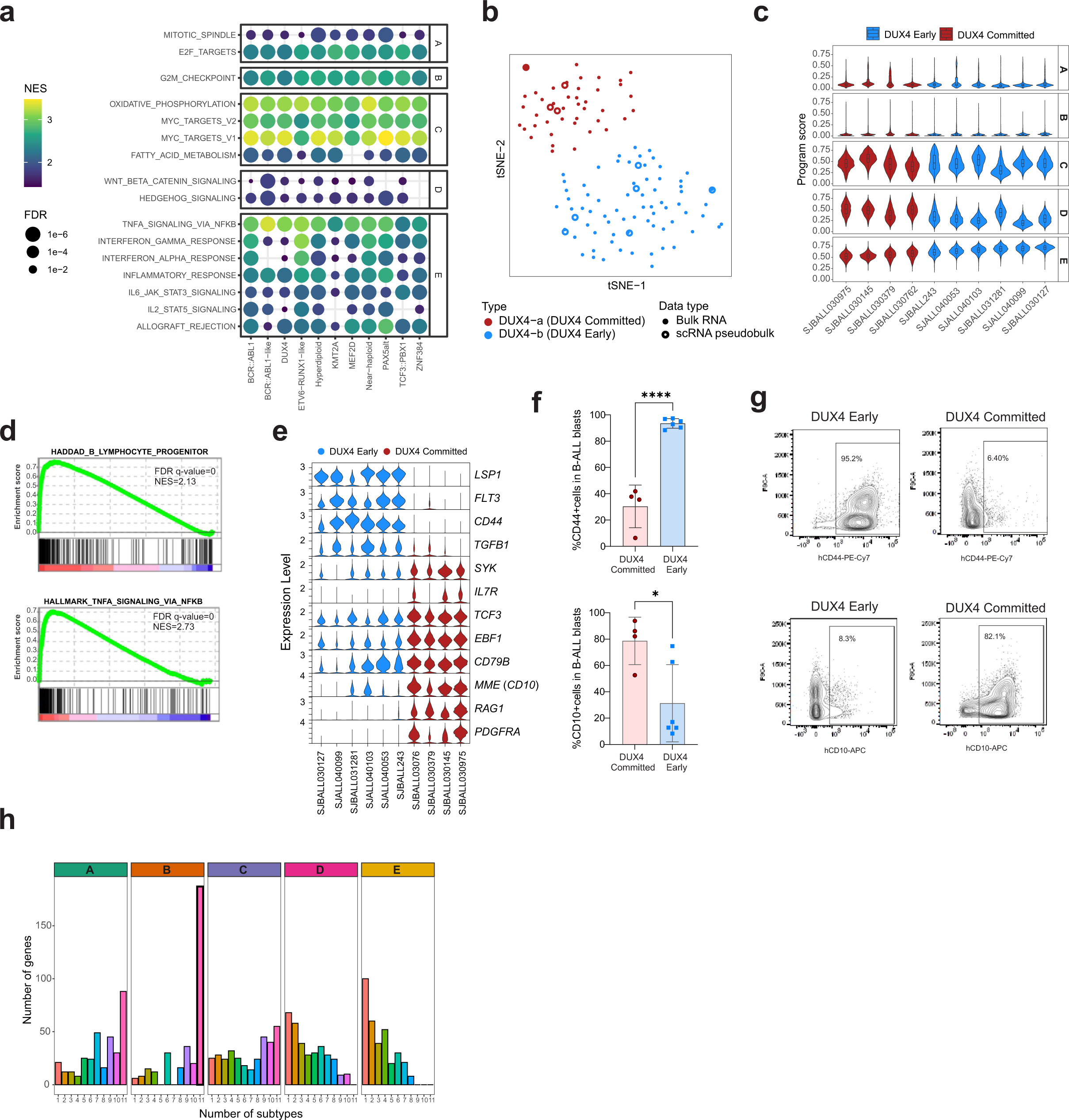
ScRNA-seq defines two subgroups for *KMT2A*-r and *BCR::ABL1* B-ALL samples. **a**) Bubble plot showing gene set enrichment analysis of the five transcriptional programs in different subtypes. Genes are ranked based on their contribution to each program in each subtype. **b**) tSNE plot showing gene expression profiling of 106 B-ALL bulk RNA-seq samples^17^ and 10 pseudobulk RNA-seq from this single-cell study. Each dot represents a sample and samples are colored by the DUX4 type (a vs b). Pseudobulk RNA-seq samples clustered with their bulk counterpart. **c**) Violin plot of the AUCell enrichment scores for the four *DUX4* subtype-level cNMF meta-programs, including A - Cell Cycle (S), B - Cell Cycle (G2/M), C - Metabolism, D - Differentiation, and E - Inflammation. **d**) Gene set enrichment analysis (GSEA) on the genes ranked by their contribution to subtype-level program 1 – Differentiation and 2 - Inflammation - in panel D identified enriched B lymphocyte progenitor pathways and TNF-alpha signaling via NFκB, respectively, in DUX4 Committed and DUX4 Early, samples. **e**) Violin plot of gene expression pattern for eight selected genes in 10 *DUX4*-r patients. **f**) Histograms showing protein expression by flow cytometric analysis of CD44 and CD10 in type a vs type b. **g**) Scatter plots showing expression of CD44 and CD10 in representative samples with DUX4 Early and DUX4 Committed. **h**) Distribution of the number of subtypes for the top 30 signature genes in the five meta programs of different subtypes indicating most of the signature genes are shared for A - Cell Cycle (S), B - Cell Cycle (G2/M), C – Metabolism but subtype-specific for D – Cell Differentiation, and E - Inflammation.

**Extended Data Figure 5.**
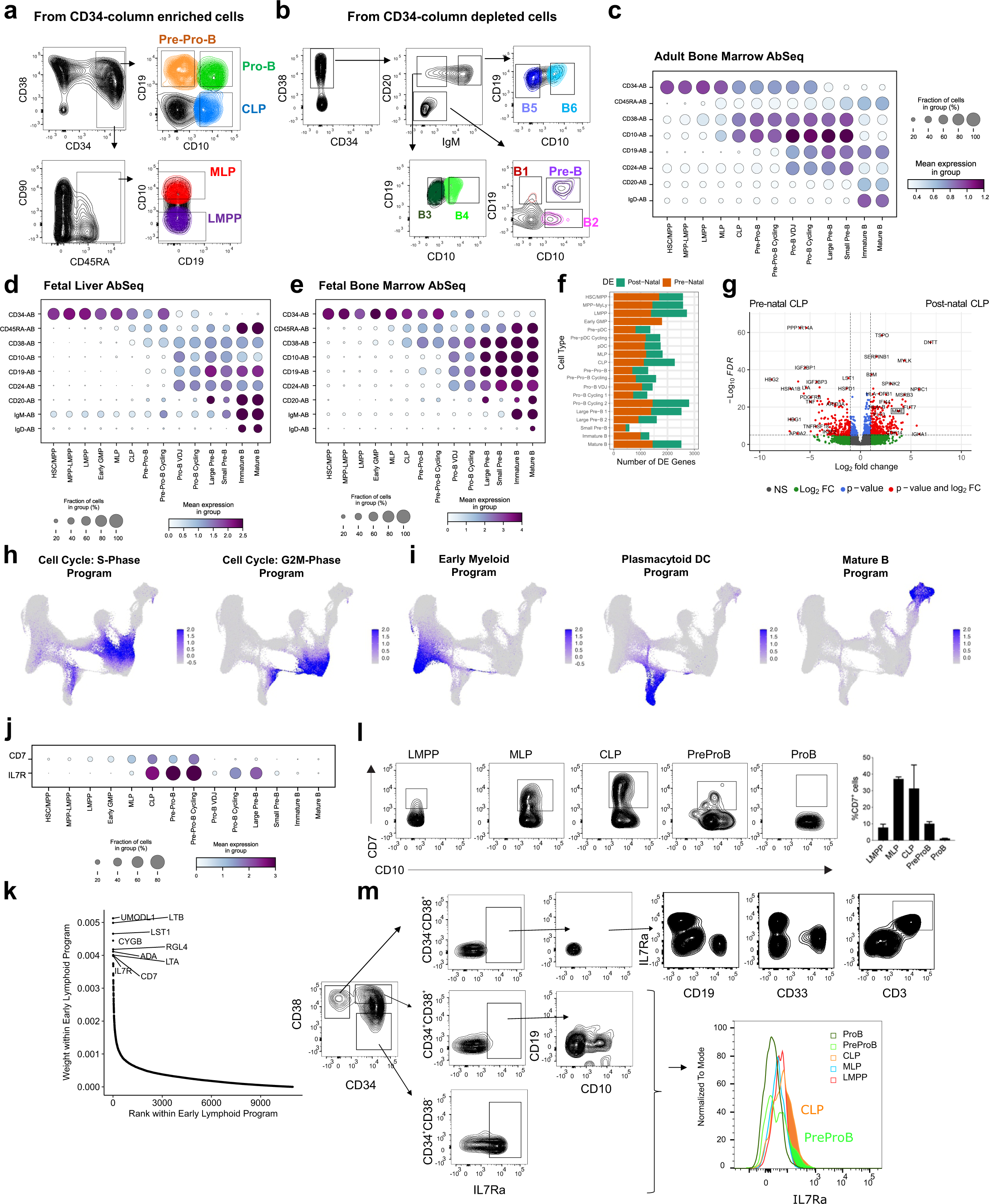
Transcriptional and functional characterization of human B cell development. **a)** Sorting scheme to identify and isolate the indicated cell populations human cord blood (hCB) enriched for CD34+ cells by MACS column (Miltenyi Biotec). **b)** Sorting scheme to identify and isolate the indicated cell populations from hCB samples depleted of CD34+ cells by MACS column. Populations from panels **a** and **b** were defined as: PreB: CD38+CD20-IgM-CD19+CD10+; B1: CD38+CD20-IgM-CD19+CD10-; B2: CD38+CD20-IgM-CD19-CD10+; B3: CD38+CD20+IgM-CD19+CD10-; B4: CD38+CD20+IgM-CD19+CD10+; B5: CD38+CD20+IgM+CD19+CD10-; B6: CD38+CD20+IgM+CD19+CD10+; **c-e)** CLR-normalized protein levels of key cell surface markers across B cell development from single cells profiled by joint scRNA + protein (AbSeq). **c)** 5,102 adult bone marrow cells profiled through BD Rhapsody from Triana et al^32^. These cell states were classified through reference map projection onto our B cell development atlas. **d)** 4,189 fetal liver cells profiled through CITE-seq from Jardine et al^30^, this data was already included within our B cell development atlas. **e)** 7,782 fetal bone marrow cells profiled through CITE-seq from Jardine et al^30^. **f)** Differential expression (DE) results between pre-natal and post-natal samples across human B-cell development. Single cells belonging to each cell state from each donor from our B cell development atlas were pooled into pseudo-bulk profiles prior to DE. Number of DE genes at FDR < 0.01 specific to either pre-natal or post-natal samples are shown. **g)** Differential expression results in CLPs from pre-natal compared to post-natal samples. *MME*, which encodes CD10, is shown within a box for emphasis. **h-i)** cNMF signature analysis in normal B-lymphoid development. Signature strength across B cell development is shown for cNMF programs corresponding to **h)** S and G2/M-phases of cell cycle and **i)** to Early Myeloid, Plasmacytoid DC, and Mature B identity. **j)** *CD7* and *IL7R* expression by cell state across the B cell development atlas. **k)** Ranking of component genes by weight within the Early Lymphoid NMF program. The top 10 genes, which include *CD7* and *IL7R*, are explicitly highlighted. **l)** Analysis of CD7 protein expression by flow cytometry in the indicated cell populations. **m)** Analysis of IL7R protein expression by flow cytometry in the indicated cell populations.

**Extended Data Figure 6.**
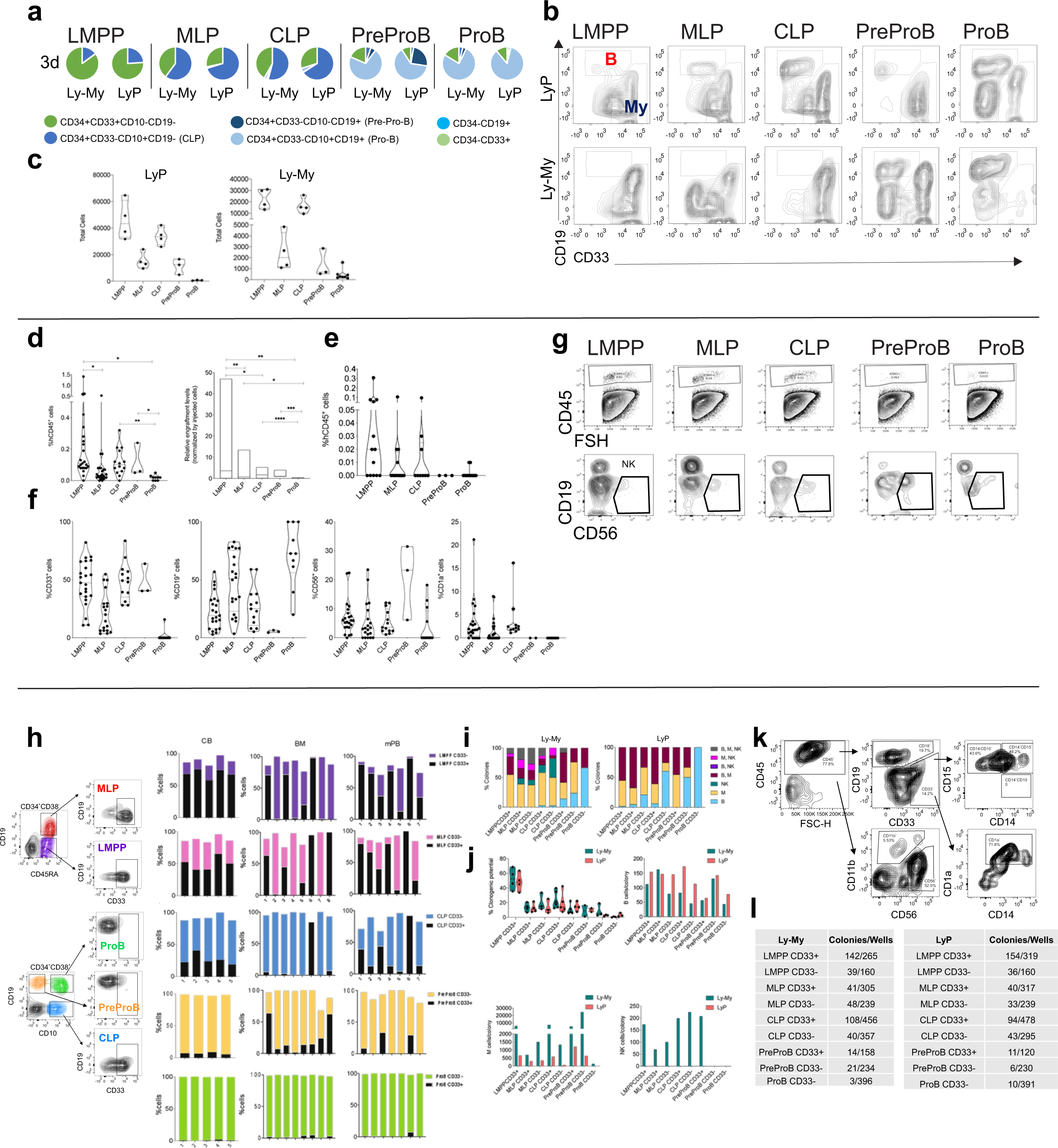
Functional modeling of flow-sorted populations. **a)** 200 cells from indicated cell populations were sorted and cultured on MS5 stroma cells for 3, 7 or 14 days (see Fig. 4 for 7 and 14 days) in Lympho-Myeloid (Ly-My) or Lymphoid (LyP) promoting medias (see Material and Methods for media composition) and analyzed by flow cytometry. The proportion of each population is represented in pie charts. **b)** Representative FACS plots from (**a**) at day 14 of culture. **c)** Total number of cells from (**a**) at day 14 of culture. **d)** Engraftment analysis of indicated cell populations after 2 weeks of interfemoral injection into NSG mice. Injected cells were: LMPP 3,000 cells/mouse; MLP: 4,000 cells/mouse; CLP: 6,000cells/mouse; Pre-ProB 6,000 cells/mouse; ProB: 10,000 cells/mouse. The percentage of human CD45+ cells and relative engraftment normalized by the number of cells injected in the right femur (injected bone) are shown. **e)** Engraftment levels in the left femur. **f)** Percentage of cells in the right femur from panel (a). **g)** Representative FACS plots from panel (**f**). **h)** Human CB, adult bone marrow (BM) and mobilized peripheral blood (MPB) were stained and analysed by flow cytometry with the indicated cell surface markers. The percentage of CD33+ and CD33-cells in each cell population is represented. CB n=5, BM n=8, MPB n=7. **i)** Single cells from indicated cell populations were sorted into MS5 stroma cells and cultured for 16-17 days with Ly-My or LyP medias. Colonies were scored under the microscope and analysed by flow cytometry for differentiation markers. The percentage of each type of colonies is shown. **j)** From (**i**) the clonogenic potential (wells with colonies/seeded wells*100) and the number of B, Myeloid or NK cells/colony are represented for both medias. **k)** Representative FACS plots from (**i**). **l)** Number of wells seeded and number of colonies for each cell population analysed.

**Extended Data Figure 7.**
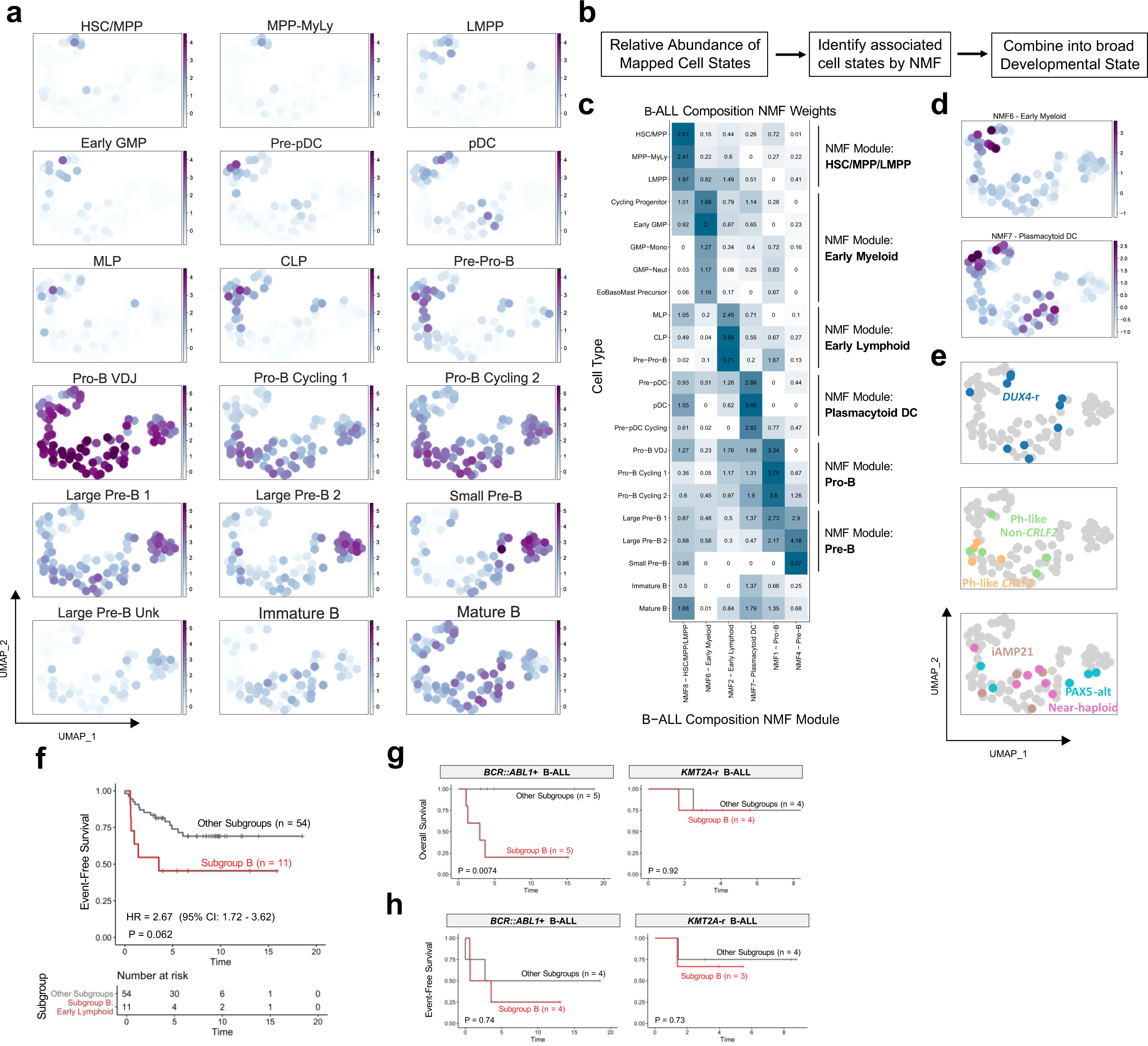
B-ALL developmental states refine existing genomic subgroups and are clinically relevant. **a**) UMAP embedding based on cell state composition from 89 B-ALL samples depicting centered log ratio (CLR) normalized abundance of each mapped cell population for each sample. **b**) Consolidation of similar B-ALL cell states into broader developmental states. To identify B-ALL states with correlated abundance across patient samples, NMF was performed on the normalized abundance scores of all mapped cell states from B-ALL bone marrow samples and NMF modules were interpreted based on the weights assigned to the cell states that comprise each module. **c**) For NMF modules that correspond to a distinct stage along B-lymphoid development, termed Developmental States, feature weights are visualized in this heatmap by constituent cell type. **d**) Abundance of broad developmental states defined by NMF spanning Myeloid Prog and Pre-pDC. **e**) Remaining genomic subtypes are overlaid based on the cell composition of their constituent patient samples. **f)** Event-free survival outcomes of patients in subgroup B compared to patients from other subgroups among samples collected from patients at initial diagnosis (n = 65). P values from likelihood ratio test are shown. **g)** Overall survival outcomes of patients in subgroup B compared to other subgroups within *BCR::ABL1*+ B-ALL (n = 10) and within *KMT2A*-r B-ALL (n = 8). P values from likelihood ratio test are shown. **h)** Event-free survival outcomes of patients in subgroup B compared to other subgroups within *BCR::ABL1*+ B-ALL (n = 8) and within *KMT2A*-r B-ALL (n = 7). P values from likelihood ratio test are shown.

**Extended Data Figure 8.**
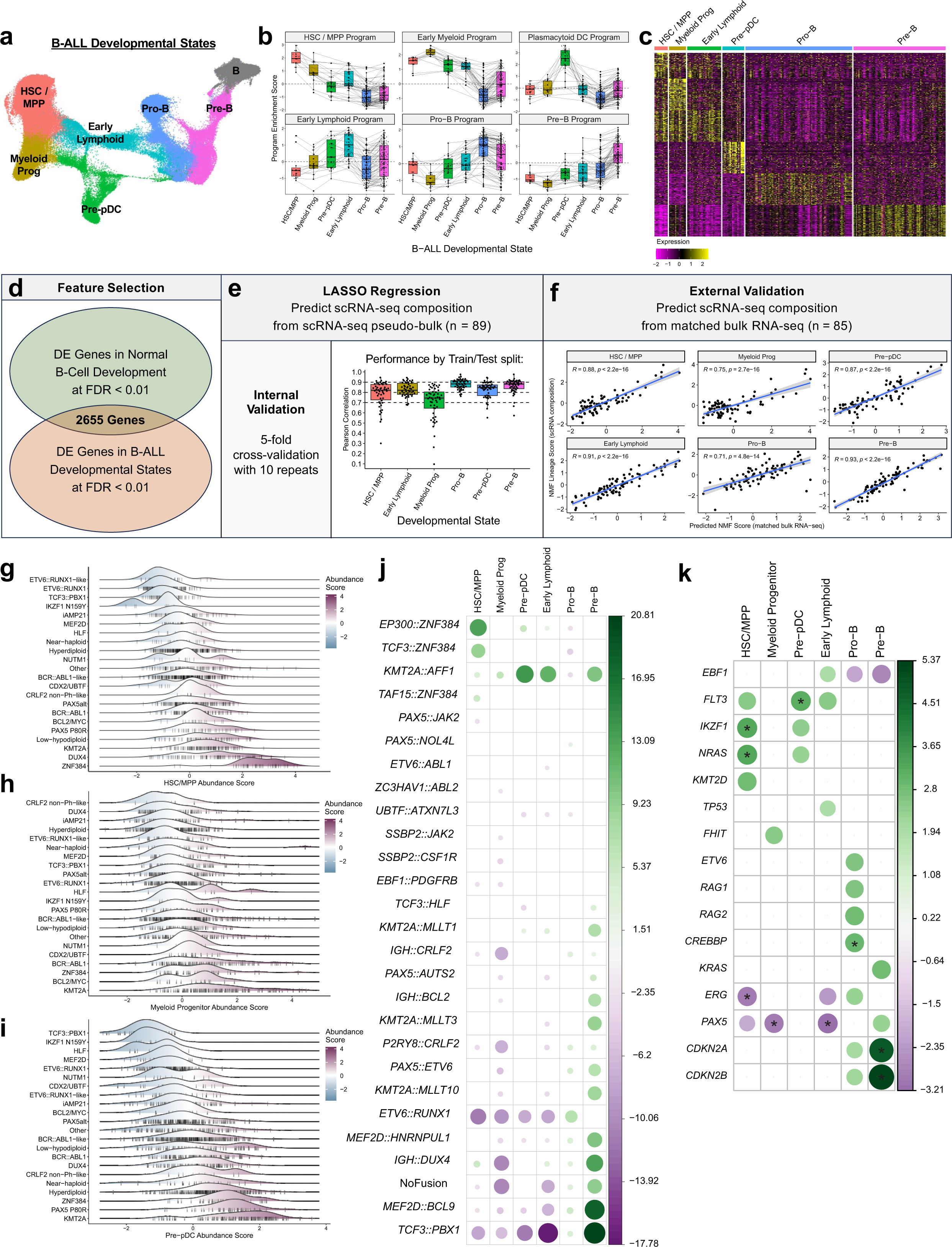
Association of B-ALL developmental states with genetic characteristics in B-ALL. **a)** Broad developmental states across B-cell differentiation implicated in B-ALL. **b)** GSVA enrichment scores for gene expression programs defined from normal B-cell development calculated in the context of B-ALL developmental states. Cells belonging each developmental state from each B-ALL sample were pooled into pseudo-bulk profiles, and developmental states from the same patient sample are connected by a line. **c)** Heatmap depicting expression of top 100 marker genes for each B-ALL developmental state. Each row corresponds to a unique marker gene while each column corresponds to a pseudo-bulk profile of each developmental state from each B-ALL sample. **d-f)** Development of gene expression scores to infer B-ALL developmental state abundance from bulk RNA-seq data. **d)** Feature selection approach prior model training. Differentially expressed genes at FDR<0.01 across both normal B-cell development as well as B-ALL developmental states were retained, yielded 2,655 genes recurrently implicated in both normal and malignant human B-cell development. **e)** LASSO regression was performed to predict developmental state abundance of 89 scRNA-seq samples. scRNA-seq profiles from each of the 89 samples were pooled into patient-level pseudo-bulk profiles to predict developmental state abundance starting from expression of the 2,655 genes. Internal cross-validation was performed to estimate the accuracy of these models in 50 train-test splits of the data, and test-set results for models corresponding to each of the six developmental states are shown. **f)** Validation of final developmental state models using matched bulk RNA-seq profiles available for 85 of the scRNA-seq samples. For each developmental state, predicted abundance from matched bulk RNA-seq data are shown in the x-axis, and the actual abundance from scRNA-seq composition data is shown on the y-axis. **g-i)** Ridge plot comparing the inferred abundance of **g)** HSC/MPP, **h)** Myeloid Progenitor, and **i)** Pre-pDC states by genomic subtype across 2,046 B-ALL patients. **j-k)** Association plots between developmental state abundance and gene fusions and mutations of B-ALL patients. The magnitude of each association, quantified as the –log10 (p value), is depicted through the size and color intensity of each dot. The direction of the association is depicted through the color, wherein higher abundance is green and lower abundance is purple. Only associations with an FDR corrected p value < 0.05 are depicted. **j)** Association between gene fusion and inferred abundance of B-ALL developmental states from 2,039 pediatric and adult patients with available fusion data. **k)** Association between genetic mutation status and inferred abundance of B-ALL developmental states from 672 pediatric patients with available mutation data. For these associations, genomic subtype was adjusted for as a covariate.

**Extended Data Figure 9.**
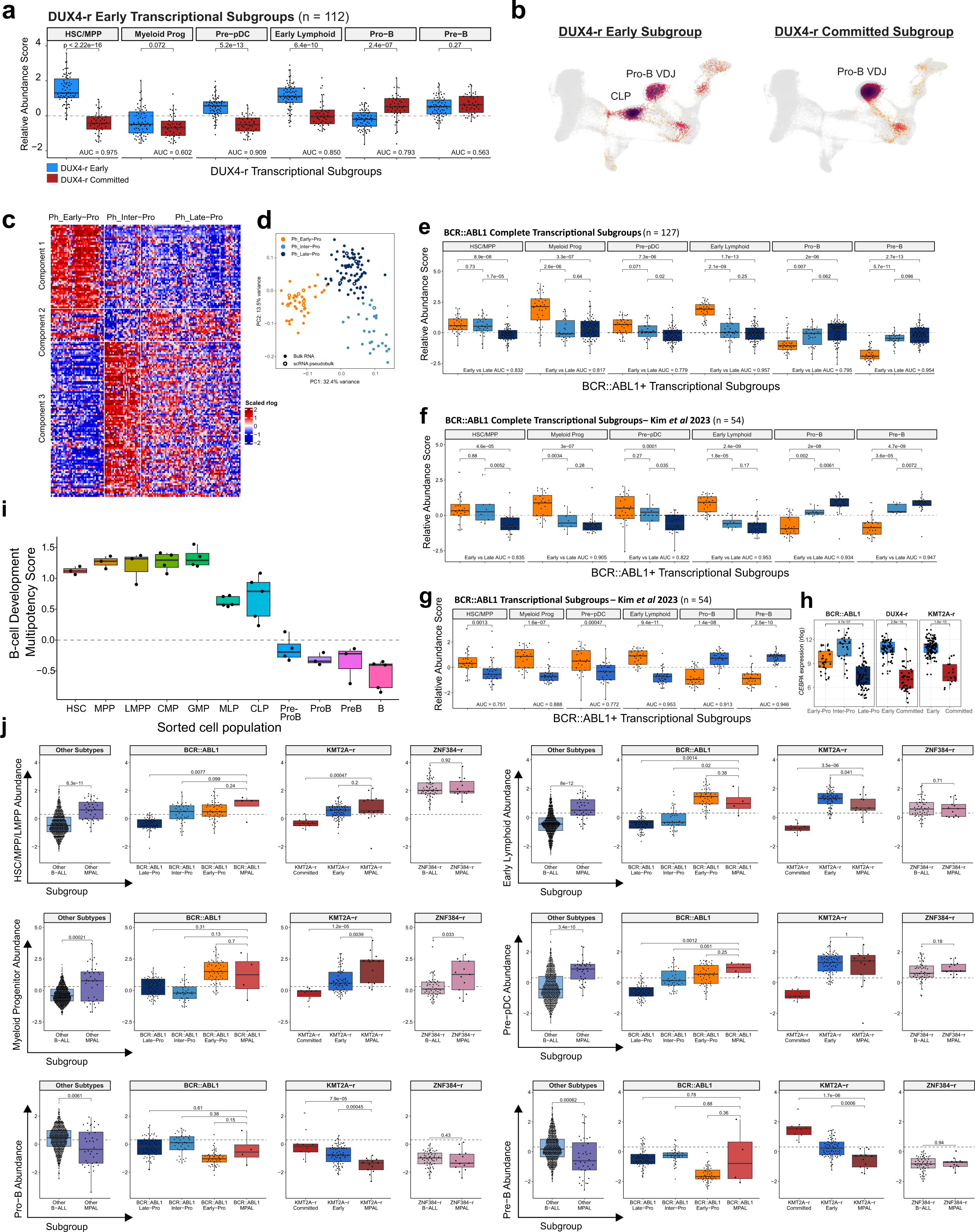
B-ALL developmental states refine existing subtypes. **a)** Developmental state abundance explains differences between Early and Committed transcriptional subsets of *DUX4-r* B-ALL. **b)** Projection results from representative patients from the Early and Committed subgroups are shown. **c**) Consensus hierarchical clustering using 163 NMF component genes Kim et al 2023 (ref^10^) resembled the three molecular subtypes of *BCR::ABL1* lymphoblastic leukemia (Ph_Early-Pro; Ph_Inter-Pro and Ph_Late-Pro). Rows represent the genes grouped into NMF components, and columns represent the 142 bulk RNA-seq samples and 11 pseudobulk scRNA-seq samples. **d**) tSNE plot showing gene expression profiling of samples in Extended Data 4E. Each dot represents a sample and samples are colored by the Ph type. Pseudobulk RNA-seq samples clustered with their bulk counterpart. **e-f)** Developmental state abundance explains differences between Early-Pro, Inter-Pro, and Late-Pro transcriptional subtypes of *BCR::ABL1* B-ALL from **e)** 127 patients within our dataset and from **f)** 54 patients from Kim et al 2023 (ref^10^). **g)** Developmental state abundance explains differences between broader Early and Committed transcriptional subsets of *BCR::ABL1* B-ALL from 54 patients from Kim et al 2023 (ref^10^). **h)** rlog normalized *CEBPA* expression across transcriptional and developmental subtypes of *BCR::ABL1* positive, *KMT2A*-r, and *DUX4*-r BALL. **i)** B-cell Multipotency scores in FACS sorted cell populations from Fig. 4. **j)** Abundance of HSC/MPP/LMPP, Early Lymphoid, Myeloid Progenitor, Pre-pDC, Pro-B and Pre-B in Early and Committed subgroups of *BCR::ABL1*, *KMT2A*-r and *ZNF384*-r with B-ALL and in “Other B-ALL”, non-*BCR::ABL1*, non-*KMT2A*-r and non-*ZNF384*-r) and in MPAL patients with same genetic drivers (*BCR::ABL1*, *KMT2A*-r and *ZNF384*-r or “Other”).

**Extended Data Figure 10.**
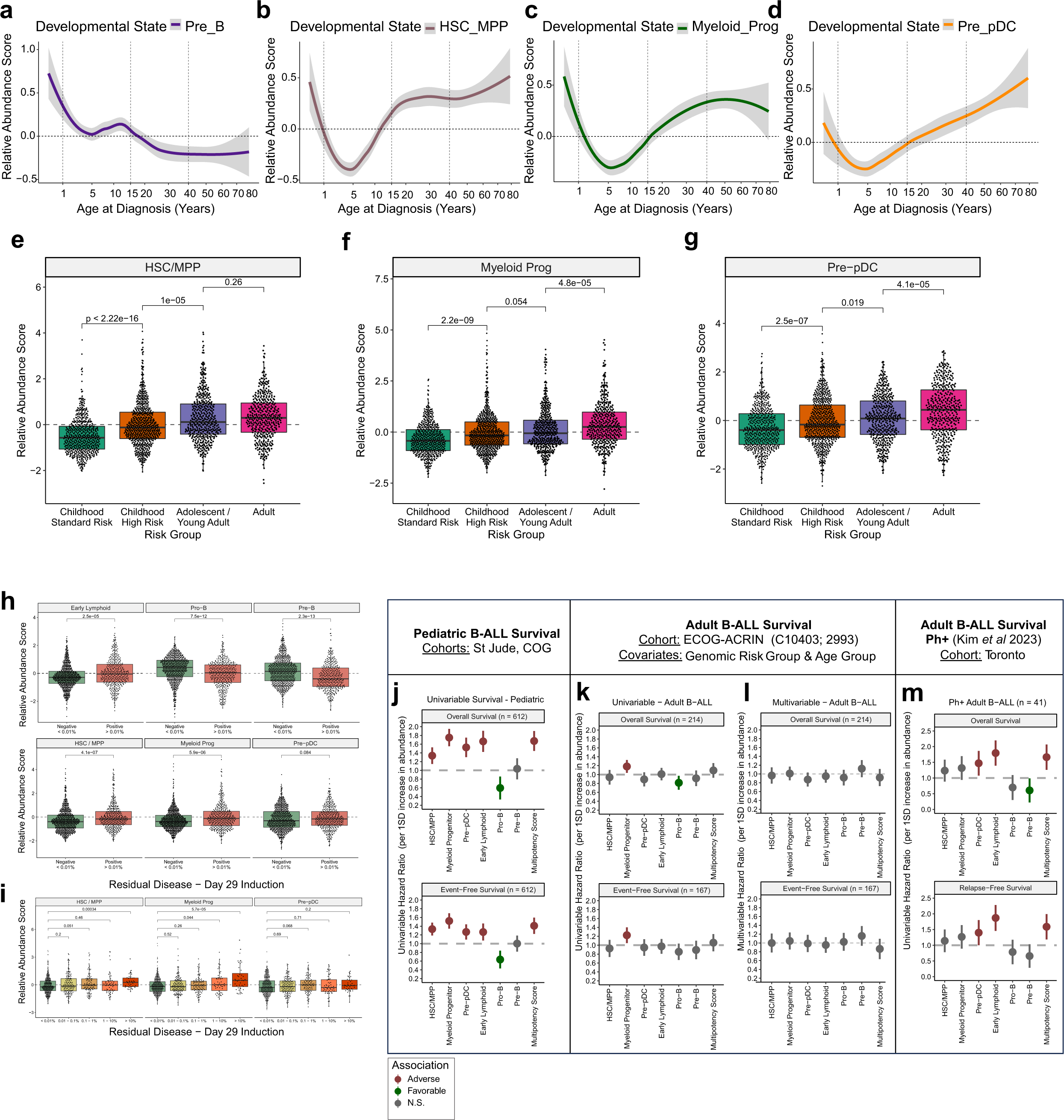
B-ALL developmental states are associated with clinical outcomes. **a-d)** Association between age at B-ALL diagnosis and inferred abundance of **a)** Pre-B, **b)** HSC/MPP, **c)** Myeloid Progenitor, and **d)** Pre-pDC developmental states. **e-g)** Association between B-ALL clinical risk groups and inferred abundance of **e)** HSC/MPP, **f)** Myeloid Progenitor, and **g)** Pre-pDC developmental states. **h)** Association between inferred developmental state at B-ALL diagnosis with measurable residual disease (MRD) status at day 29 of induction chemotherapy administration across 1192 pediatric B-ALL patients for which MRD information was available. **i)** Association between inferred abundance of HSC/MPP, Myeloid Progenitor, and Pre-pDC states at B-ALL diagnosis with residual disease levels at day 29 of induction chemotherapy administration for 792 pediatric B-ALL patients for which residual disease levels were available. **j)** Association of B-ALL developmental states with survival outcomes within 826 pediatric B-ALL patients. Hazard ratios from univariate cox regression with overall survival and event-free survival are reported for each standard deviation increase in inferred abundance. **k-l)** Association of B-ALL developmental states with survival outcomes within 214 genetically diverse adult B-ALL patients. Hazard ratios for overall survival and event-free survival from **k)** univariate cox regression or **l)** multivariable Cox regression accounting for age group and genomic subtype are shown are reported for each standard deviation increase in inferred abundance. **m)** Association of B-ALL developmental states with survival outcomes within 41 *BCR::ABL1* B-ALL patients from Kim et al^10^. Hazard ratios from univariate Cox regression with overall survival and event-free survival are reported for each standard deviation increase in inferred abundance.

## ONLINE METHODS

### Cohort description

We analyzed children, adolescents and young adults (AYA) with newly diagnosed B-ALL (N=80) and matched diagnosis-relapse B-ALL (N=5). All patients were enrolled-on St. Jude Children’s Research Hospital sponsored trials, including Total Therapy XV (ClinicalTrials.gov Identifier NCT00137111), Total Therapy XVI (NCT00549848) and Total Therapy XVII (NCT03117751).. Detailed clinical information for each case is provided in **Supplementary Table 1**. The study was conducted in accordance with the Declaration of Helsinki and approved by the Institutional Review Board. Written informed consent was obtained from the patient, parents or guardians and assent from the patients, as appropriate.

### Single-cell RNA sequencing

Frozen mononuclear cells from diagnosis and relapse bone marrow samples were thawed, counted and up to 2 million cells were subjected to dead cell removal (Miltenyi Biotec, catalogue number 130-090-101) following manufacturer’s instructions. Isolated live cells were washed three times with 1X Phosphate-Buffered Saline (PBS, calcium and magnesium free) containing 0.04% weight/volume BSA (Thermo Fisher Scientific, catalogue number AM2616) and automatedly counted by the Countess 3 Automated Cell Counter (Thermo Fisher Scientific) and manually by the Neubauer hemocytometer. From each sample, we calculated the volume of cells to load to have a desired recovery target of 8,000-10,000 cells and loaded on Chromium Next GEM Chip K (10X Genomics; PN-2000182) with Master Mix from Chromium Next GEM Single Cell 5’ Reagent Kits v2 (Dual Index) (10X Genomics; PN-1000263), Single Cell VDJ 5’ Gel Beads (10X Genomics; PN-1000264) and Partitioning Oil (10X Genomics; PN-2000190) following standard manufacturer’s protocols. Briefly, nanoliter-scale gel beads-in-emulsion (GEMs) were generated, lysed and 10x barcoded, full-length cDNA from polyadenylated mRNA was produced. Next, GEMs were broken and pooled fractions were recovered. Dynabeads MyOne SILANE (10X Genomics; PN-2000048) were used to purify the first-strand cDNA from the post GEM–RT reaction mixture. Barcoded, full-length cDNA was amplified via PCR, purified by SPRIselect Reagent (Beckman Coulter, B23318) and quantified using High Sensitivity D5000 ScreenTape (5067-5592) with High Sensitivity D5000 Reagents (5067-5593) at the Agilent 2200 TapeStation system. Enzymatic fragmentation and size selection were used to optimize the cDNA amplicon size. To construct the final libraries, P5, P7, i5 and i7 sample indexes, and Illumina R2 sequence (read 2 primer sequence) were added via End Repair, A-tailing, Adaptor Ligation, and Sample Index PCR (10X Genomics; Library Construction Kit, 16 reactions PN-1000190 and Dual Index Kit TT Set A, 96 reactions PN-1000215). Final library quality was assessed using High Sensitivity D1000 ScreenTape (5067-5584) with High Sensitivity D1000 Reagents (5067-5585) at the Agilent 2200 TapeStation system. Illumina-ready dual index libraries were sequenced at the recommended depth and aiming at 50,000 reads/cell at the Hartwell Center at SJCRH on the Illumina NovaSeq according to manufacturer’s recommendations.

### Single-cell RNA sequencing analysis

Single-cell RNA-seq data were aligned and quantified using the Cell Ranger (v5.0.1) pipeline (http://www.10xgenomics.com) against the human genome GRCh38 (refdata-gex-GRCh38-2020-A). Cells with mitochondria content >8%, or number of detected genes <500, or total number of detected molecules <2500 were removed using Seurat (4.0.6, https://satijalab.org/seurat)^49^ using Uniform Manifold Approximation and Projection (UMAP) reduction and characterized based on gene expression of major haemopoietic cell types. Doublets and multiplets were filtered out using DoubletFinder with the core statistical parameters (nExp and pK) determined automatically using recommended settings^50^. Ambient cell free mRNA contamination was estimated using SoupX^51^. Copy number variations were inferred from single cell gene expression using inferCNV with parameters HMM=TRUE, analysis_mode = "subclusters", k_obs_groups = 8, denoise=TRUE (v.1.11.1, https://github.com/broadinstitute/inferCNV). The groups with similar copy number pattern were then merged. Differential expression analysis between different clusters of cells were performed using FindMarkers function from Seurat package.

### Single-cell whole genome sequencing (scWGS)

ScWGS was performed as previously described^52^. Briefly, frozen bone marrow mononuclear cells from SJE2A063, SJE2A066 and SJE2A067 were thawed and stained with Alexa Fluor® 700 Mouse Anti-Human CD45 (BD Biosciences, #560566) and PE Mouse Anti-Human CD3 (BD Biosciences, #561809) and single cell fluorescence-activated cell sorting (FACS) in 96-well plate for CD3-CD45dim (leukemic blasts) and CD3+CD45bright (normal T cells). Sorted single-cell were processed by PicoPLEX Gold Single Cell DNA-Seq Kit (Takara Bio USA, catalogue number R300670), according to manufacturer’s instructions. Briefly, single cells were lysed, DNA was pre-amplified by linear amplification, purified by AMPure XP (Beckman Coulter, catalogue number A63880) and exponentially amplified and indexed with Illumina primers from HT Unique Dual Index Kit (1–96) (Takara Bio USA, catalogue number 634752). Libraries were purified by AMPure XP and quality was checked by Agilent D1000 ScreenTape 5067-5582) with D1000 Reagents (5067-5583) prior sequencing.

### scWGS analysis

scWGS data were mapped to human reference genome GRCh37 by BWA (version 0.7.12). Samtools (version 1.5) was used to estimate the sequencing depth for each segment inferred from bulk WGS data of SJE2A063 and SJE2A067. To generate the coverage normalization index, segment coverages of individual normal cell were firstly divided by the median segment coverage across the genome, and then the median value of each genomic segment among all the normal cells was used as the normalization index. To estimate the copy number for individual tumor cells, segment coverages were first divided by the coverage normalization index, then by the median segment coverage across the genome, and multiplied by two due to diploidy.

### Non-negative matrix factorization (NMF)

To determine the transcriptional programs within leukemic cells, we performed NMF using cNMF (v1.1)^24^ for the largest inferred copy number clone of each sample (clone C1 in Fig. 2a). For this analysis we used the top 2,000 overdispersed genes found in the upper 50% of malignant cells exhibiting high total RNA counts . To identify the most stable and accurate number of factors (k) for each sample, 100 iterations of NMF with different random seeds were run for values of k from 5 to 10. For each value of k, the Silhouette score, measuring the stability of the components, and the Frobenius error were computed, and the k maximizing the Silhouette score and minimizing the Frobenius error was selected for each sample, as implemented in cNMF. For each of the resulting factors, we considered the 30 genes with the highest NMF scores to be characteristic of that factor. For each subtype, hierarchical clustering of the scores for each program was used to identify main correlated sets of programs. The 30 genes with the highest average NMF score within each correlated program set (excluding ribosomal protein genes) were then used to define subgroup-specific metaprograms. Each program was annotated using pathway enrichment analysis.

### Human cord blood (CB) samples

Human CB samples were obtained with informed consent from Trillium Health, Credit Valley and William Osler Hospitals according to procedures approved by the University Health Network (UHN) Research Ethics Board. Mononuclear cells (MNC) from pools of male and female CB units (∼4-15 units) were obtained by centrifugation on Lymphoprep medium (Stem Cell Technologies), and after ammonium chloride lysis MNC were enriched for CD34+ cells by positive selection with the CD34 Microbead kit (Miltenyi Biotec, Cat#130-097-047) and LS column purification with MACS magnet technology (Miltenyi Biotec, Cat#130-042-401). Resulting CD34+ CB cells were viably stored in 50% PBS, 40% fetal bovine serum (FBS) and 10% DMSO at -80°C or −150°C.

### Flow Cytometric Analysis and Sorting

For cell sorting, CD34+ and CD34-human CB cells were thawed via slow dropwise addition of X-VIVO 10 medium with 50% fetal bovine serum (FBS) and DNaseI (200 μg/ml). Cells were spun at 350g for 10 minutes (min) at 4°C and then resuspended in phosphate-buffered saline (PBS) + 2.5% FBS. For all in vitro and in vivo experiments, the full stem and progenitor hierarchy sort was performed as described in Notta et al.^53^ and shown in Fig. 4 and **Extended Data Figs. 5a-b**. Cells were resuspended in 100 μl per 1x10^6^ cells and stained in two subsequent rounds for 15 min at room temperature each. See **Supplementary Table 17** for full list and details of the antibodies used.

### Cell Culture

For in vitro experiments sorted cells were cultured in 96 well-plate round bottom with the indicated cell media. Lympho-Myeloid media (Ly-My): H5100 media supplemented with 5% FCS, 1x L-Glutamine (Thermo Fisher), 1x Penicillin-Streptomycin (Thermo Fisher) and the following cytokines: SCF (100ng/mL), TPO (50ng/mL), IL7 (20ng/mL), IL2 (10ng/mL) and Flt3 (10ng/mL). Lymphoid promoting media (LyP): IMDM media supplemented with 10% Bit, 1x L-Glutamine (Thermo Fisher), 1x Penicillin-Streptomycin (Thermo Fisher) and the following cytokines: SCF (100ng/mL), TPO (50ng/mL), IL7 (20ng/mL).

### Single cell Stromal Assays

Single cell in vitro assays were set up as described previously^54^ with low passage murine MS-5 stroma cells seeded at a density of 1,500 cells per 96-well and grown for 2-4 days in H5100 media (Stem Cell Technologies). One-day prior to coculture initiation, the H5100 media was removed and replaced with 100 μl erythro-myeloid differentiation media: StemPro-34 SFM media (Thermo Fisher) with the supplied supplement, 1x L-Glutamine (Thermo Fisher), 1x Penicillin-Streptomycin (Thermo Fisher), 0.02% Human LDL (Stem Cell Technologies) and the following cytokines: FLT3L (20 ng/mL), GM-CSF (20 ng/mL), SCF (100 ng/mL), TPO (100 ng/mL), EPO (3 U/mL), IL-2 (10 ng/mL), IL-3 (10 ng/mL), IL-6 (50 ng/mL), IL-7 (20 ng/mL), and IL-11 (50 ng/mL). Sorted single-cells were deposited into the MS-5 seeded 96-well plates (80 wells/96-well plate) using the FACSAria II (BD). Colonies were scored after 15-17 days under the microscope and every individual well containing a visible colony was stained with antibodies listed in **Supplementary Table 17** and analyzed by flow cytometry using a BD FACSCelesta instrument equipped with a high throughput sampler (HTS).

### Mouse experiments

Mouse experiments were done in accordance with institutional guidelines approved by University Health Network Animal care committee. All in vivo experiments were done with 8- to 12-week-old female/male *NOD.Cg-Prkdc^scid^Il2rg^tm1Wjl^/SzJ* (NSG) mice (JAX) mice that were irradiated the day before intrafemural cell injection. All mice were housed at the animal facility (ARC) at Princess Margaret Cancer Centre in a room designated only for immunocompromised mice with individually ventilated racks equipped with complete sterile micro-isolator caging (IVC), on corn-cob bedding and supplied with environmental enrichment in the form of a red house/tube and a cotton nestlet. Cages are changed every <7 days under a biological safety cabinet. Health status is monitored using a combination of soiled bedding sentinels and environmental monitoring.

### Xenotransplantation

The progeny of 10,000 CD34+CD38-transduced cells one day after transduction were intra-femoral injected in aged and gender matched 8–12-week-old male and female recipient NSG. At indicated time points, mice were euthanized, injected femur and other long bones (non-injected femur, tibiae) were flushed separately in Iscove’s modified Dulbecco’s medium (IMDM) (Thermo Fisher) + 5%FBS and 5% of cells were analysed for human chimerism along with antibodies listed in **Supplementary Table 17**. Sick and mis-injected mice were excluded from analysis.

### RNA-seq processing and analysis

Freshly sorted populations from 3-5 independent CB pools were directly resuspended and frozen (-80°C) in PicoPure RNA Isolation Kit Extraction Buffer (ThermoFisher, Cat#KIT0204). RNA was isolated using the PicoPure RNA Isolation Kit (Thermo Fisher, Cat#KIT0204) according to manufacturer’s instructions. Samples that passed quality control according to integrity (RIN>8) and concentration as verified on a Bioanalyzer pico chip (Agilent Technologies) were subjected to further processing by the Center for Applied Genomics, Sick Kids Hospital: cDNA conversion was performed using SMART-Seq v4 Ultra Low Input RNA Kit for Sequencing (Takara) and libraries were prepared using Nextera XT DNA Library Preparation Kit (Illumina). Equimolar quantities of libraries were pooled and sequenced with 4 cDNA libraries per lane on a High Throughput Run Mode Flowcell with v4 sequencing chemistry on the Illumina 30 HiSeq 2500 following manufacturer’s protocol generating paired-end reads of 125-bp in length to reach depth of 55-75 million reads per sample. Reads were then aligned with STAR v2.5.2b against hg38 and annotated with ensembl v90. Default parameters were used except for the following: chimSegmentMin 12; chimJunctionOverhangMin 12; alignSJDBoverhangMin 10; alignMatesGapMax 100000; alignIntronMax 100000; chimSegmentReadGapMax parameter 3; alignSJstitchMismatchNmax 5 -1 5 5. Read counts were generated using HTSeq v0.7.2 and general statistics were obtained from picard v2.6.0. Differential gene expression was performed using edgeR_3.24.3 following recommended practices.

### ATAC-seq processing and analysis

Library preparation for ATAC-Seq was performed on 1000-5000 cells with Nextera DNA Sample Preparation kit (Illumina), according to previously reported protocol^55^. 4 ATAC-seq libraries were sequenced per lane in HiSeq 2500 System (Illumina) to generate paired-end 50-bp reads. Reads were mapped to hg38 using BWA (0.7.15) using default parameters. Duplicate reads, reads mapped to mitochondria, an ENCODE blacklisted region or an unspecified contig were removed (Encode Project Consortium, 2012). MACS (2.2.5) was used to call peaks in mapped reads. A catalogue of all peaks was obtained by concatenating all peaks and merging any overlapping peaks. Peaks were considered unique to one condition or another if they were present in at least 2 out of three replicates but not in the contrasting condition. Homer (4.11.1) was used to calculate enrichment of subsets of peaks using default parameters plus the catalogue of called peaks as background.

### Cross-ontogeny reference map of B cell development

In order to increase the resolution of our normal B-lymphoid reference and expand the source of the reference to include fetal tissue which are hypothesized to be a potential origin of B-ALL, we expanded our reference map by incorporating scRNA-seq data from additional studies incorporating fetal liver^29^, fetal bone marrow^30^, umbilical cord blood (Human Cell Atlas CB), and a fourth study incorporating progenitors across ontology^31^.

Briefly, scRNA-seq data from each study was mapped to the bone marrow reference map and any cells positioned within the B-lymphoid lineage (HSC, MPP, LMPP, MLP, CLP (Early Pro-B), Pre-Pro-B, Pro-B, Pre-B, Immature B, Mature B) were retained as well as cells mapping to proximal states branching off of B-lymphoid development (early GMP, Pre-pDC, Pre-pDC Cycling, pDC).

As a result, scRNA-seq data spanning ontogeny (fetal liver, fetal BM, cord blood, pediatric BM, adult BM) from 90 donors across eight studies representing B-cell development and proximal branch points (GMP, pDC) were integrated to develop a comprehensive map of B-cell development. The full list of studies, tissues and technologies represented in the B-cell developmental map is provided in the **Supplementary Table 9.** See **Supplementary Note 1** in the **Supplementary Information** file for the fully detailed method description.

### Composition analysis of B-ALL data

#### BALL scRNA-seq Mapping

For mapping of B-ALL scRNA-seq samples, filtered counts from 10x were loaded into Seurat (v4.3.0) and subject to further QC with the following filters: nFeature_RNA >500, nCount_RNA >2500, and pct.mito <8. Filtered single-cell transcriptomes were first projected onto a complete Bone Marrow reference map (BoneMarrowMap v0.1.0, https://github.com/andygxzeng/BoneMarrowMap) based on 2,386 highly variable genes spanning human bone marrow cells. Cells that were assigned to the following cell state by BoneMarrowMap (HSC, MPP-MyLy, LMPP, Early GMP, MLP, MLP-II, Pre-pDC, Pre-pDC Cycling, pDC, CLP, Pre-ProB, Pro-B VDJ, Pro-B Cycling, Large Pre-B, Small Pre-B, Immature B, Mature B) were retained. These cells were subsequently mapped onto our B cell development map using symphony (v0.1.1), based on 950 highly variable genes spanning human B cell development.

#### BALL scRNA-seq Composition Analysis

For composition analysis of B-ALL, the number of mapped single-cell transcriptomes that passed a mapping QC filter was counted for each sample. Cell states with less than 100 cells present across all samples were filtered out. These frequency statistics for each cell state within each sample were used.

To cluster patient samples by stages of B cell development, 21 cell states spanning B cell development were retained. Cell counts across these 21 cell states for each patient were subject to CLR normalization with multiplicative replacement using the package scikit-bio (v0.5.6). Dimensionality reduction and clustering was performed using scanpy (v1.9.1). Briefly, a neighbourhood graph was constructed based on cosine similarity across the n = 6 top PCs, incorporating n = 10 nearest neighbours. UMAP reduction was performed at min.dist = 0.2 and leiden clustering at a resolution = 1.2. This resulted in seven patient subgroups on the basis of cell composition across B cell development.

#### NMF Developmental State Analysis

To simplify downstream analyses, we sought to bundle specific cell states into broader categories based on correlated abundance across patient samples. To do so, cell counts across 55 cell states for each patient were subject to Centered Log ratio (CLR) normalization with multiplicative replacement using the package skbio, and NMF analysis was performed on this normalized composition data. Ten NMF components, each corresponding to groups of cell states that vary across samples. Out of these ten components, six correspond to discrete stages of B cell development, hereafter Developmental States. These comprise of Pro-B (NMF1), Early Lymphoid (NMF2), Pre-B (NMF4), Myeloid Progenitor (NMF6), Pre-pDC (NMF7), and HSC/MPP/LMPP (NMF8).

### Quantification in B-ALL RNA-seq

The relative abundance of each developmental state is represented by the score for each corresponding NMF component within each patient. To estimate the abundance of each developmental state in bulk RNA-seq data, we identified biologically relevant genes which were correlated to the abundance of each developmental state by integrating three approaches of feature selection. See **Supplementary Note 2** in the **Supplementary Information** file for the fully detailed method description.

### Statistical Analysis in B-ALL Cohorts

#### Inferring Developmental State Abundance in bulk RNA-seq

Briefly, the abundance of each developmental state is represented by a gene expression score, comprising a linear equation with coefficients/weights assigned by LASSO for each constituent gene. These equations are then applied to normalized bulk RNA-seq data from B-ALL patients wherein the normalized expression of each gene is multiplied by the appropriate co-efficient, and the sum of these products is used and standardized across patients. Each standardized score represents the relative abundance of a developmental state within an individual patient. Gene expression scores were calculated on *vst*-normalized^56^ data from bulk RNA-seq of 2,046 B-ALL patients.

#### Association with genomic characteristics of B-ALL

Gene expression scores were calculated on *vst*-normalized^56^ data from bulk RNA-seq of 2,046 B-ALL patients. Associations between developmental state and genomic characteristics were evaluated through a generalized linear model using glmnet (v4.1.3) wherein inferred abundance of each developmental state was used as the dependent variable and genomic category was used as the independent variable, stratifying on originating institute. For associations between mutations and developmental state, genomic subtype was used as a covariate. The resulting t-statistic and FDR corrected significance was visualized through package corrplot (v0.84). Unless otherwise indicated, all comparisons between groups were performed through a Wilcoxon rank sum test.

#### Association with patient survival outcomes

Survival associations were determined through cox proportional hazards regression using package survival (v2.44.1.1) with either overall or event-free survival outcomes as the dependent variable and inferred developmental state abundance as the independent variable. In pediatric B-ALL, survival analysis was performed across subsets of patients with available RNA-seq data from St Jude’s and Children’s Oncology Group (COG) cohort, in adult B-ALL survival analysis was performed within a subset of patients with available RNA-seq data from the European Co-operative Oncology Group (ECOG) cohort. Multivariable analysis in pediatric and adult patients was performed with Clinical Risk Group (Childhood SR, Childhood HR, AYA, Adult) and Genomic Subtype Risk Group (Favorable, Intermediate, and Unfavorable, Unclassified) as covariates. Hazard ratios are reported for each standard deviation increase in inferred abundance of a developmental state. Significance was calculated through nested likelihood ratio test (LRT) wherein the performance of a model with Clinical Risk + Genomic Subtype Risk + Developmental State Abundance is compared to the performance of a baseline model with only Clinical Risk + Genomic Subtype Risk information. This analysis was repeated independently for each developmental state.

Genomic Subtype Risk Group assignments were determined based on the genomic subtype and differed for adult and pediatric B-ALL. Among pediatric B-ALL, samples were classified as Favorable (*ETV6::RUNX1*, Hyperdiploid, *DUX4-*rearranged, iAMP21, *NUTM1-*rearranged, *PAX5* P80R, and *ZNF384-*rearranged), Intermediate (*PAX5*alt, *BCR::ABL1*-like other, *TCF3::PBX1*, Near-haploid, *ETV6::RUNX1*-like), Unfavorable (*BCR::ABL1*, *BCR::ABL1*-like *CRLF2-* rearranged*, HLF-*rearranged, *KMT2A-*rearranged, low hypodiploid, *MEF2D, BCL2/MYC)*, or unclassified (B-Other, *CRLF2* non-*BCR::ABL1*-like, *KMT2A*-like, *ZNF384*-like, *ZEB2/CEBP*, *IKZF1* N159Y). Among adult B-ALL, samples were classified as Favorable (*ETV6::RUNX1*, *ETV6::RUNX1*-like, Hyperdiploid, *DUX4*, *PAX5* P80R, *TCF3::PBX1*, and *ZNF384*), Intermediate (*PAX5*alt, *MEF2D*), Unfavorable (*BCL2/MYC*, *KMT2A*, low hypodiploid, near haploid, *BCR::ABL1*-like *CRLF2-*rearranged, and *BCR::ABL1*-like other), and Unclassified (*BCR::ABL1*, KMT2A-like, ZNF384-like, B-Other, *CRLF2* non-*BCR::ABL1*-like, *CDX2/UBTF*, *HLF*, iAMP21, *IKZF1* N159Y).

#### Association with ex vivo drug response

Gene expression profiles of 595 B-ALL samples from Lee *et al* ^42^ were used to link developmental state abundance with *ex vivo* sensitivity to 18 therapeutic agents. Gene expression scores were calculated using logTPM normalized gene expression data obtained from the publication to infer abundance of each developmental state. Pearson correlation and FDR-corrected significance of the association between abundance of each developmental state and the area under the dose-response curve (AUC), wherein lower AUC values denote higher sensitivity. Pearson values are multiplied by -1 such that positive values indicate that higher abundance of a developmental state is associated with higher *ex vivo* sensitivity to a drug.

#### Comparison between B-ALL and MPAL

Gene expression profiles of 1194 B-ALL samples and 125 MPAL samples for which sequencing data was uniformly pre-processed was obtained from Montefiori *et al*^41^. Gene expression scores were calculated using *rlog*-normalized gene expression data obtained from the publication to infer abundance of each developmental state. For comparisons of developmental state abundance, T/Myeloid lineage MPALs were removed from the analysis to focus on B/Myeloid lineage MPALs and Acute Undifferentiated Leukemias (AULs).

#### Re-analysis of BCR::ABL1 transcriptional subgroups

Gene expression profiles of 57 *BCR::ABL1* lymphoblastic leukemia samples were *vst*-normalized and gene expression scores were calculated to infer abundance of each developmental state. Transcriptional subgroup labels of Early-Pro, Inter-Pro, and Late-Pro were obtained from the original study. Survival associations were performed on 41 adult samples with outcomes data (overall survival and relapse-free survival) using cox proportional hazards regression with inferred developmental state abundance as the independent variable. Hazard ratios are reported for each standard deviation increase in inferred abundance of a developmental state. This analysis was repeated independently for each developmental state.

## REFERENCES

1. Arber, D.A., et al. International Consensus Classification of Myeloid Neoplasms and Acute Leukemias: integrating morphologic, clinical, and genomic data. Blood 140, 1200–1228 (2022).

2. Alaggio, R., et al. The 5th edition of the World Health Organization Classification of Haematolymphoid Tumours: Lymphoid Neoplasms. Leukemia 36, 1720–1748 (2022).

3. Moorman, A.V. New and emerging prognostic and predictive genetic biomarkers in B-cell precursor acute lymphoblastic leukemia. Haematologica 101, 407–416 (2016).

4. Hunger, S.P. & Mullighan, C.G. Acute Lymphoblastic Leukemia in Children. N Engl J Med 373, 1541–1552 (2015).

5. Duncavage, E.J., et al. Genomic profiling for clinical decision making in myeloid neoplasms and acute leukemia. Blood 140, 2228–2247 (2022).

6. Stanulla, M., et al. IKZF1(plus) Defines a New Minimal Residual Disease-Dependent Very-Poor Prognostic Profile in Pediatric B-Cell Precursor Acute Lymphoblastic Leukemia. J Clin Oncol 36, 1240–1249 (2018).

7. Iacobucci, I., Kimura, S. & Mullighan, C.G. Biologic and Therapeutic Implications of Genomic Alterations in Acute Lymphoblastic Leukemia. J Clin Med 10(2021).

8. Coustan-Smith, E., et al. New markers for minimal residual disease detection in acute lymphoblastic leukemia. Blood 117, 6267–6276 (2011).

9. Duffield, A.S., Mullighan, C.G. & Borowitz, M.J. International Consensus Classification of acute lymphoblastic leukemia/lymphoma. Virchows Arch 482, 11–26 (2011).

10. Kim, J.C., et al. Transcriptomic classes of BCR-ABL1 lymphoblastic leukemia. Nat Genet 55, 1186–1197 (2023).

11. Caron, M., et al. Single-cell analysis of childhood leukemia reveals a link between developmental states and ribosomal protein expression as a source of intra-individual heterogeneity. Sci Rep 10, 8079 (2020).

12. Turati, V.A., et al. Chemotherapy induces canalization of cell state in childhood B-cell precursor acute lymphoblastic leukemia. Nat Cancer 2, 835–852 (2021).

13. Witkowski, M.T., et al. Extensive Remodeling of the Immune Microenvironment in B Cell Acute Lymphoblastic Leukemia. Cancer Cell 37, 867–882 e812 (2020).

14. Wu, L., et al. Single-Cell Transcriptome Analysis Identifies Ligand-Receptor Pairs Associated With BCP-ALL Prognosis. Front Oncol 11, 639013 (2021).

15. Anand, P., et al. Single-cell RNA-seq reveals developmental plasticity with coexisting oncogenic states and immune evasion programs in ETP-ALL. Blood 137, 2463–2480 (2020).

16. Wang, X., et al. Single-Cell RNA-Seq of T Cells in B-ALL Patients Reveals an Exhausted Subset with Remarkable Heterogeneity. Adv Sci (Weinh) 8, e2101447 (2021).

17. O’Byrne, S., et al. Discovery of a CD10-negative B-progenitor in human fetal life identifies unique ontogeny-related developmental programs. Blood 134, 1059–1071 (2019).

18. Jackson, T.R., Ling, R.E. & Roy, A. The Origin of B-cells: Human Fetal B Cell Development and Implications for the Pathogenesis of Childhood Acute Lymphoblastic Leukemia. Front Immunol 12, 637975 (2021).

19. Zeng, A.G.X., et al. A cellular hierarchy framework for understanding heterogeneity and predicting drug response in acute myeloid leukemia. Nat Med 28, 1212–1223 (2022).

20. van Galen, P., et al. Single-Cell RNA-Seq Reveals AML Hierarchies Relevant to Disease Progression and Immunity. Cell 176, 1265–1281 e1224 (2019).

21. Patel, A.P., et al. Single-cell RNA-seq highlights intratumoral heterogeneity in primary glioblastoma. Science 344, 1396–1401 (2019).

22. Gu, Z., et al. PAX5-driven subtypes of B-progenitor acute lymphoblastic leukemia. Nat Genet 51, 296–307 (2019).

23. Brady, S.W., et al. The genomic landscape of pediatric acute lymphoblastic leukemia. Nat Genet 54, 1376–1389 (2022).

24. Kotliar, D., et al. Identifying gene expression programs of cell-type identity and cellular activity with single-cell RNA-Seq. Elife 8(2019).

25. Lee, R.D., et al. Single-cell analysis identifies dynamic gene expression networks that govern B cell development and transformation. Nat Commun 12, 6843 (2021).

26. Oetjen, K.A., et al. Human bone marrow assessment by single-cell RNA sequencing, mass cytometry, and flow cytometry. JCI Insight 3(2018).

27. Ainciburu, M., et al. Uncovering perturbations in human hematopoiesis associated with healthy aging and myeloid malignancies at single-cell resolution. Elife 12(2023).

28. Setty, M., et al. Characterization of cell fate probabilities in single-cell data with Palantir. Nat Biotechnol 37, 451–460 (2019).

29. Popescu, D.M., et al. Decoding human fetal liver haematopoiesis. Nature 574, 365–371 (2021).

30. Jardine, L., et al. Blood and immune development in human fetal bone marrow and Down syndrome. Nature 598, 327–331 (2021).

31. Roy, A., et al. Transitions in lineage specification and gene regulatory networks in hematopoietic stem/progenitor cells over human development. Cell Rep 36, 109698 (2021).

32. Triana, S., et al. Single-cell proteo-genomic reference maps of the hematopoietic system enable the purification and massive profiling of precisely defined cell states. Nat Immunol 22, 1577–1589 (2021).

33. Kondo, M., Weissman, I.L. & Akashi, K. Identification of clonogenic common lymphoid progenitors in mouse bone marrow. Cell 91, 661–672 (1997).

34. Hao, Q.L., et al. Identification of a novel, human multilymphoid progenitor in cord blood. Blood 97, 3683–3690 (2001).

35. Van de Sande, B., et al. A scalable SCENIC workflow for single-cell gene regulatory network analysis. Nat Protoc 15, 2247–2276 (2020).

36. Doulatov, S., et al. Revised map of the human progenitor hierarchy shows the origin of macrophages and dendritic cells in early lymphoid development. Nat Immunol 11, 585–593 (2010).

37. Alexander, T.B., et al. The genetic basis and cell of origin of mixed phenotype acute leukaemia. Nature 562, 373–379 (2010).

38. Dickerson, K.M., et al. ZNF384 Fusion Oncoproteins Drive Lineage Aberrancy in Acute Leukemia. Blood Cancer Discov 3, 240–263 (2018).

39. Khabirova, E., et al. Single-cell transcriptomics reveals a distinct developmental state of KMT2A-rearranged infant B-cell acute lymphoblastic leukemia. Nat Med 28, 743–751 (2022).

40. Chen, C., et al. Single-cell multiomics reveals increased plasticity, resistant populations, and stem-cell-like blasts in KMT2A-rearranged leukemia. Blood 139, 2198–2211 (2021).

41. Montefiori, L.E., et al. Enhancer Hijacking Drives Oncogenic BCL11B Expression in Lineage-Ambiguous Stem Cell Leukemia. Cancer Discov 11, 2846–2867 (2021).

42. Lee, S.H.R., et al. Pharmacotypes across the genomic landscape of pediatric acute lymphoblastic leukemia and impact on treatment response. Nat Med 29, 170–179 (2023).

43. Mehtonen, J., et al. Single cell characterization of B-lymphoid differentiation and leukemic cell states during chemotherapy in ETV6-RUNX1-positive pediatric leukemia identifies drug-targetable transcription factor activities. Genome Med 12, 99 (2020).

44. Lee, B.J., et al. CD19-directed immunotherapy use in KMT2A-rearranged acute leukemia: A case report and literature review of increased lymphoid to myeloid lineage switch. Am J Hematol 97, E439–E443 (2022).

45. Tirtakusuma, R., et al. Epigenetic regulator genes direct lineage switching in MLL/AF4 leukemia. Blood 140, 1875–1890 (2022).

46. Haddox, C.L., et al. Blinatumomab-induced lineage switch of B-ALL with t(4:11)(q21;q23) KMT2A/AFF1 into an aggressive AML: pre- and post-switch phenotypic, cytogenetic and molecular analysis. Blood Cancer J 7, e607 (2017).

47. Iacobucci, I. & Mullighan, C.G. KMT2A-rearranged leukemia: the shapeshifter. Blood 140, 1833–1835 (2022).

48. Novakova, M., et al. DUX4r, ZNF384r and PAX5-P80R mutated B-cell precursor acute lymphoblastic leukemia frequently undergo monocytic switch. Haematologica 106, 2066–2075 (2021).

49. Hao, Y., et al. Integrated analysis of multimodal single-cell data. Cell 184, 3573–3587 e3529 (2021).

50. McGinnis, C.S., Murrow, L.M. & Gartner, Z.J. DoubletFinder: Doublet Detection in Single-Cell RNA Sequencing Data Using Artificial Nearest Neighbors. Cell Syst 8, 329–337 e324 (2019).

51. Young, M.D. & Behjati, S. SoupX removes ambient RNA contamination from droplet-based single-cell RNA sequencing data. Gigascience 9(2020).

52. Gao, Q., et al. The genomic landscape of acute lymphoblastic leukemia with intrachromosomal amplification of chromosome 21. Blood (2023).

53. Notta, F., et al. Distinct routes of lineage development reshape the human blood hierarchy across ontogeny. Science 351, aab2116 (2016).

54. Wagenblast, E., et al. Functional profiling of single CRISPR/Cas9-edited human long-term hematopoietic stem cells. Nat Commun 10, 4730 (2019).

55. Buenrostro, J.D., Giresi, P.G., Zaba, L.C., Chang, H.Y. & Greenleaf, W.J. Transposition of native chromatin for fast and sensitive epigenomic profiling of open chromatin, DNA- binding proteins and nucleosome position. Nat Methods 10, 1213–1218 (2013).

56. Love, M.I., Huber, W. & Anders, S. Moderated estimation of fold change and dispersion for RNA-seq data with DESeq2. Genome Biol 15, 550 (2014).

