## Supplementary Information for "SINGLE CELL DISSECTION OF DEVELOPMENTAL ORIGINS AND TRANSCRIPTIONAL HETEROGENEITY IN B-CELL ACUTE LYMPHOBLASTIC LEUKEMIA"

### Supplementary Tables

**Supplementary Table 1.**
Clinical and genetic features of B-ALL samples analyzed by scRNA-seq.

**Supplementary Table 2.**
Quality control metrics for the 89 scRNA-seq data.

**Supplementary Table 3.**
Subtypes and library protocols for the 2,046 bulk RNA-seq data.

**Supplementary Table 4.**
Number of blast cells, number of copy number subclones and number of cells in each subclone based on inferCNV copy number analysis.

**Supplementary Table 5.**
Copy number and cluster of each segment in each cell from scWGS-seq of samples SJE2A063_D and SJE2A067_D.

**Supplementary Table 6.**
Expression pattern of genes on q arm of chromosome 1 in different subclones of sample SJE2A063_D.

**Supplementary Table 7.**
The 30 genes with the highest NMF score for each meta-program in each subtype from sample-level NMF analysis.

**Supplementary Table 8.**
The 30 genes with the highest NMF score for each meta-program in subtypes *DUX4*-r from subtype-level NMF analysis.

**Supplementary Table 9.**
Details of studies, tissues, and technologies represented in B-cell development map.

**Supplementary Table 10.**
Top 100 genes driving each NMF gene expression program in normal B cell development. Genes were ranked by NMF co-efficient weight and the top 100 genes for each program is shown, these can be used as gene sets for scoring program activity in bulk and single cell datasets.

**Supplementary Table 11.**
Cell type composition data from B-ALL scRNA-seq samples. For each B-ALL sample, the abundance of each cell type is shown as the raw number of cells, the proportion of all cells from that sample, and the centered log ratio (CLR) normalized abundance within that sample.

**Supplementary Table 12.**
B-ALL developmental state abundance within scRNA-seq data. For each B-ALL sample, the relative abundance of each broad developmental state is reported as the value of the corresponding NMF component derived from cell type composition analysis.

**Supplementary Table 13.**
Top 100 marker genes for each B-ALL developmental state based on differential expression analysis across 89 B-ALL scRNA-seq samples. These can be used as gene sets for scoring in single cell datasets.

**Supplementary Table 14.**
Model weights for predicting B-ALL developmental state abundance from normalized bulk RNA-seq profiles. For each B-ALL developmental state as well as the B-ALL multipotency score, the LASSO co-efficient for each gene within the model is reported.

**Supplementary Table 15.**
Normalized abundance of each cell type for the 2,046 bulk RNA-seq samples.

**Supplementary Table 16.**
Non-malignant cell type composition for the 89 scRNA-seq samples.

**Supplementary Table 17.**
List of antibodies used for sorting of distinct populations along B-cell development.

### Supplementary Results

#### Clonal genetic heterogeneity

To investigate clonal genetic heterogeneity within leukemic blasts we used inferCNV. Most of the samples (N=46, 51.7%) had one single predominant CNV clone, however among the remaining samples, 16.9% (N=24) had two clones, 3.4% (N=15) had three clones and less than 1.5% had 4 or more clones (4 clones in 3 samples; 8 clones in 1 sample) (**Fig. 2a** and **Supplementary Table 4**). The lower limit of subclone detection was 10 cells. In samples with more than one copy number clone, the predominant clone accounted for over 50% of cells, while the additional clones were often subclonal (**Fig. 2a**). Near haploid samples showed chromosome losses in all leukemic cells but not in normal cells suggesting that these gross chromosomal abnormalities occur earlier most likely in one single event. Minor clones showed partial loss of chromosomal regions. For example, in SJHYPO117, all leukemic cells showed loss of chromosomes 1, 2, 3, 5, 6, 7, 9, 10, 11, 12, 13, 15, 16, 17, 19, 20 and 22, suggesting that these alterations were earlier events arising in the founder clone. However, three clones were depicted by inferCNV and differentiated in the inferred copy number state from scRNA-seq of chromosomes 8, 14 and 18. Chromosomes 8 and 18 were wild-type in both clone 1 (18.2% of blasts) and clone 2 (58.5% of blasts) but loss in clone 3 (23.3% of blasts), while we observed a gradual loss of chromosome 14, which was retained in clone 1, partially lost in clone 2 and completely lost in clone 3 (**Figs. 2b-c**). Interestingly, the inferCNV clones were associated with distinct gene expression profiles. While clone 1 and 2 clustered together, clone 3 with extra chromosomal losses showed a distinct cluster in the UMAP visualization (**Fig. 2d**). All the remaining near haploid samples showed loss of entire chromosomes in all cells, suggesting that each chromosome was lost in a single event and not through gradual losses (**Fig. 2b**).

We thereafter analyzed inferCNV in samples with hyperdiploidy (N=10) (**Extended Data Figs. 2a-c**) B-ALL[1,2]. In all samples we observed chromosomes gains in all cells consistent with a common origin rather than gradual accumulation of alterations as previously demonstrated[3]. However, in a subset of cases a second subclone was observed involving a gain of new additional chromosomes (e.g., gain of chromosome 12 in SJALL040100 in less than 1.5% of blasts; chromosome 17 in SJBALL030276 in less than 1% of blasts and chromosome 18 in SJBALL030285 in 12% of cells) or extra gain of amplified chromosomes (e.g., in SJBALL030491 with extra gain of chromosome X and in SJBALL030821 with extra gain of chromosome 6) (**Extended Data Figs. 2a-c**).

**Characterization of human B cell development**

Prior studies have noted the existence of a fetal-specific population along B-cell development, namely a CD10-CD19- Early Lymphoid Progenitor (ELP)[4-6]. Notably, we found that fetal and post-natal tissues shared common transcriptional states spanning B cell development, however we did observe some variation in surface marker expression by ontogeny. In post-natal tissue, CD10 upregulation occurs at the MLP stage (**Extended Data Fig. 5c**) while in fetal tissue CD10 upregulation is delayed until the Pro-B stage (**Extended Data Figs. 5d-e**). Differential expression analysis revealed transcriptional differences between pre-natal and post-natal tissue sources that persist across B cell differentiation and confirmed transcriptional upregulation of the CD10-encoding transcript *MME* within post-natal CLP (**Extended Data Figs. 5f-g**). Thus, despite differences in surface marker expression, pre-natal “ELPs” represent an equivalent cellular state to post-natal CLPs and both will hereafter be referred to as CLPs.

**Composition of non-malignant cells in B-ALL**

The 89 B-ALL samples processed by scRNA-seq were not subjected to blast enrichment prior processing and thus they included both leukemic and normal hematopoietic cells in a ratio proportional to the blast cell count. Since the presence of normal hematopoietic cells offers the opportunity to explore cell microenvironment composition and its correlation with molecular subtypes, we analyzed the type and proportion of non-blast cells (HSPCs, monocytes, erythroid cells, DCs, B cells, T/NK cells and plasmablasts) within each sample and across B-ALL subtypes (**Supplementary Fig. 1a** and **Supplementary Table 16**) and found enrichment of specific normal cell types according to the B-ALL molecular subtype. For example, we observed a significantly higher proportion of non-classical monocytes (CD14dim, CD16+) in samples with *ETV6::RUNX1*-like and *TCF3::PBX1* (one-way ANOVA, $p < 0.0001$) (**Supplementary Fig. 1b**), confirming previous scRNAseq data from *ETV6::RUNX1* samples[7]. To investigate subtype-specific expression patterns of this cell population, we performed differential gene expression analysis between CD16 positive monocytes in *ETV6::RUNX1*-like and CD16 positive cells in all the other B-ALL subtypes. This analysis showed a distinct expression signature in CD16+ cells from *ETV6::RUNX1*-like, with overexpression of genes involved in WNT (*LBH*), PI3K/AKT (*AKT3*) or interleukin signaling (*CSF1R*). Moreover, GSEA revealed enrichment of RHO GTPASE signaling and downregulation of ribosomal proteins in normal cells from *ETV6::RUNX1*-like compared to those from other subtypes (**Supplementary Fig. 1c**). The presence of non-classical monocytes in *ETV6::RUNX1*-like extends previous data in *ETV6::RUNX1*-positive B-ALL[7].

Interestingly, dendritic cells (overexpression of *IL3RA* and *CST3*) were instead enriched in *BCR::ABL1* positive ALL and specifically they were only found in samples with an early pre-B differentiation stage (Ph Early)(**Supplementary Fig. 1d-e**).

Overall, these findings suggest a dynamic crosstalk between leukemia cells and their microenvironment.

### Supplementary Notes

#### Supplementary Note 1: B-Development Map Construction

In order to increase the resolution of our normal B-lymphoid reference and expand the source of the reference to include fetal tissue which are hypothesized to be a potential origin of B-ALL, we expanded the BoneMarrowMap resource (https://github.com/andygxzeng/BoneMarrowMap) by incorporating scRNA-seq data from additional studies incorporating fetal liver[8], fetal bone marrow[9], umbilical cord blood (HCA cord blood), and a fourth study incorporating progenitors across ontology[10].

Briefly, scRNA-seq data from each additional study was filtered at nGenes > 500, nGenes < 6000, and pct.mito < 8, and was subsequently mapped to the bone marrow reference map and cells assigned to the B-lymphoid lineage (HSC, MPP-MyLy, LMPP, MLP, MLP-II, CLP, Pre-Pro-B, Pro-B VDJ, Pro-B Cycling, Large Pre-B, Small Pre-B, Immature B, Mature B) were retained alongside cells assigned to proximal cell states branching off of B-lymphoid development (Early GMP, Pre-pDC, Pre-pDC Cycling, pDC).

Among single cell transcriptomes positioned within B cell development and proximal lineages, scRNA-seq data spanning ontogeny (fetal liver, fetal BM, cord blood, pediatric BM, adult BM) from 90 donors across eight studies representing B-cell development and proximal branch points (GMP, pDC) were integrated to develop a comprehensive map of B-cell development including a total of 130,085 single cell transcriptomes.

This map of B cell development comprised of single-cell transcriptomes from seven studies: Human Cell Atlas, n = 15,689; Oetjen *et al* 2018 (ref[11]), n = 6,593; Ainciburu *et al* 2023 (ref[12]), n = 23,278; Setty *et al* 2019 (ref[13]), n = 11,071; Popescu *et al* 2019 (ref[8]), n = 5,471; Jardine *et al* 2021 (ref[9]), n = 42,877; Roy *et al* 2021 (ref[10]), n = 25,106. Transcriptomes from Granja *et al* 2019 (ref[14]) and Mende *et al* 2022 (ref[15]) were excluded due to high rates of gene dropout particularly involving B cell receptor genes. This atlas also spanned 90 donors across five tissues (fetal liver, n = 20,944; fetal bone marrow, n = 39,680; cord blood, n = 5,680; pediatric bone marrow, n = 5,870; adult bone marrow, n = 57,911), and three technologies (10x 3’ V2, n = 74,990; 10x 3’ V3, n = 41,576; 10x 5’, n = 13,519).

**Highly variable gene selection**

Proper highly variable gene (HVG) selection was critical in minimizing batch effects from technology and donor. Our final HVG selection approach divided the cohort by sequencing technology and used the sc.pp.highly_variable_genes function within scanpy (v1.9.1) adjusting for donor ID among cells sequenced with each technology. HVGs surpassing a normalized dispersion threshold > 1 within each technology (10x 3’ V2, n = 425 genes; 10x 3’ V3, n = 606 genes; 10x 5’, n = 581 genes) were retained and the union of these genes across all three technologies was used. This resulted in 950 final variable genes that were highly variable across B cell development, irrespective of donor, within any of the three technologies. These 950 genes were used for subsequent dimensionality reduction.

**Dimensionality Reduction and Clustering**

Dimensionality reduction and cell type annotation was performed in line with BoneMarrowMap (https://github.com/andygxzeng/BoneMarrowMap) wherein an iterative grid search was performed across multiple batch correction parameters and multiple dimensionality reduction parameters to identify a UMAP embedding that exhibited a continuous hematopoietic differentiation manifold while retaining adequate distance between non-cycling CD34+ Pro-B and CD34- Pre-B cells.

Parameters for the final embedding included 950 HVGs as described above, batch correction with harmony (v0.1.1) across 90 individual donors with parameters theta = 1, max.iter.cluster = 150, and max.iter.harmony = 20. A neighbourhood similarity matrix was constructed for the top

100 neighbours of each cell by cosine distance across the top 30 harmony components and UMAP reduction was performed with min.dist = 0.29 and spread = 0.9.

To identify cell states within the data, leiden clustering was performed using scanpy (v1.9.1) at resolutions ranging from 0.5 to 20. To guide cluster selection, we performed SingleR (v1.6.1) scoring with bulk RNA-seq profiles from cord blood sorted fractions spanning human B cell development, as well as bulk RNA-seq from fetal sorted fractions[16]. Scores for consecutive populations were contrasted with one another (e.g. HSC vs LMPP; Pro-B vs Pre-B). Clusters were chosen to reflect optimal cut-offs pertaining to SingleR cell fraction scores, known marker gene expression, and BoneMarrowMap annotations. Briefly, broad classifications from low resolution leiden clustering would be refined into specific cell states with higher resolution leiden clustering to align cell state assignments with marker expression and SingleR scores. Any remaining unlabeled cell types were labeled through KNN classification based on harmony components. These assignments were subsequently validated by marker gene expression, enrichment of SingleR bulk RNA-seq fraction scores, BoneMarrowMap assignments, and projection of Abseq data[17].

**Consensus Non-negative Matrix Factorization**

Consensus non-negative matrix factorization[18] was run with 100 iterations on the single cells to pull out transcriptional gene programs of variation in an unsupervised manner. Raw counts were fed into the –counts flag, and SCTransform V2 (ref[19]) corrected counts specifically from the data slot of the SCT assay were fed into the –tpm flag of the cNMF command line function with 10 workers. Following running cNMF across various components, the optimal number of components was selected by assessing the silhouette score (stability) and Frobenius reconstruction error (error), as implemented in cNMF. The transcriptomic signatures were annotated by cell type enrichment and an overrepresentation analysis with the function, 'fora' from the package fgsea[20] on the top 100 genes from each signature with pathways from MSigDb. The feature genes were ranked by the z-score regression coefficients.

**pySCENIC Regulon Inference**

The single cells of the B cell development reference map were downsampled to the metacell level using the divide and conquer algorithm from the Metacell2 package[21]. The 950 HVGs used to construct the reference map were used as input into the algorithm guiding feature gene selection, thus optimizing clustering for metacell partitioning. The approach generated 6,394 metacells with a target metacell size of 75,000 UMI. Gene regulatory network inference was performed using a candidate list of TFs[22], to identify co-expression modules using pySCENIC[23], particularly on the raw count metacells, to reduce runtime. Candidate regulons were pruned using the annotations of transcription factor motifs, 'motifs-v9-nr.hgnc-m0.001-o0.0.tbl.' Subsequently, Cistarget was employed using the 'mc9nr' databases, which include known human TF motifs annotated at: a) 500 bp upstream and 100 bp downstream of the transcriptional start site (TSS) and b) 10 kb centered around the TSS. Log-transformed counts of the Metacells were used as input for CisTarget, and drop-out masking was applied. Finally, the transcription factor regulons were scored using AUCell in the single-cell dataset, quantifying transcription factor activity.

### Supplementary Note 2: Quantification in B-ALL RNA-seq

The relative abundance of each developmental state is represented by the score for each corresponding NMF component within each patient. To estimate the abundance of each developmental state in bulk RNA-seq data, we identified biologically relevant genes which were correlated to the abundance of each developmental state by integrating three approaches of feature selection.

**First**, we created patient-level pseudo-bulk profiles from the 89 patient samples and identified genes that were significantly associated with abundance of each developmental state. Briefly, pseudo-bulk samples were *vst* normalized using DESeq2 (v1.32.0) and a Pearson test was performed between normalized gene expression and abundance of each developmental state across 89 patient samples. Filtering at FDR < 0.05 resulted in thousands of significantly associated genes for each developmental state, thus further filtering was required. First, adaptive K1 thresholding, adapted from the AUCell R package, was performed within significantly positively correlated genes and within significantly negatively correlated genes.

**Second**, we identified differentially expressed genes across normal B cell development. Cells belonging to each normal cell type within a healthy donor from the B cell development reference map were pooled into pseudo-bulk profiles. Only profiles comprised of ≥ 5 cells were retained. One-vs-all differential expression was performed between each cell state and all others using DESeq2 (v1.32.0), correcting for Study ID and Tissue Source (Fetal Liver, Fetal Bone Marrow, Cord Blood, Bone Marrow) as covariates. Among the correlated genes, only those that were differentially expressed across normal B cell development at FDR < 0.01 were retained.

**Third**, we identified differentially expressed genes across B-ALL developmental states. Cells belonging to each developmental state within a B-ALL patient were pooled into pseudo-bulk profiles. Only profiles comprised of ≥ 100 cells were retained. One-vs-all differential expression was performed between each developmental state and all others using DESeq2 (v1.32.0), correcting for Patient ID as a covariate. Among the correlated genes, only those that were differentially expressed across B-ALL developmental states at FDR < 0.01 were retained.

**Together**, for each developmental state only the genes that met the following criteria were used for model training:

1) Positively or Negatively correlated with abundance of this B-ALL developmental state at FDR < 0.05 and exceeding an adaptive positive or negative threshold

2) Significantly differentially expressed across normal B cell development

3) Significantly differentially expressed across B-ALL developmental states

Using genes that met these criteria (**Extended Data Figure 8d**), LASSO regression was performed on developmental state abundance using *vst* normalized gene expression data from pseudo-bulk profiles of 89 B-ALL patient samples, using R package glmnet (v4.1.2). Briefly, to optimize model parameters 5-fold cross-validation was performed within each dataset with 10 repeats. Data was shuffled in between repeats, resulting in performance estimates for 50 random train/test splits (**Extended Data Figure 8e**). Pearson correlations across each train/test split was used as a measure of accuracy on unseen data.

Final models for each developmental state were trained using LASSO, with leave-one-out cross-validation to determine the lambda value corresponding to the lowest root mean square prediction error (RMSE). To further reduce the number of features in the model, the largest lambda within one standard error of the lowest RMSE was used for the training the final model. This resulted in a different gene expression model for inferring abundance of each B-ALL developmental state, with all six models captured within 145 genes.

To validate these gene expression scores, we obtained matched bulk RNA-seq data for 85 out of the 89 B-ALL samples profiled by scRNA-seq. Inferred Developmental State abundance calculated from bulk RNA-seq profiles were compared against actual Developmental State abundance derived from the original scRNA-seq composition analysis (**Extended Data Figure 8f**).

### Supplementary Figure

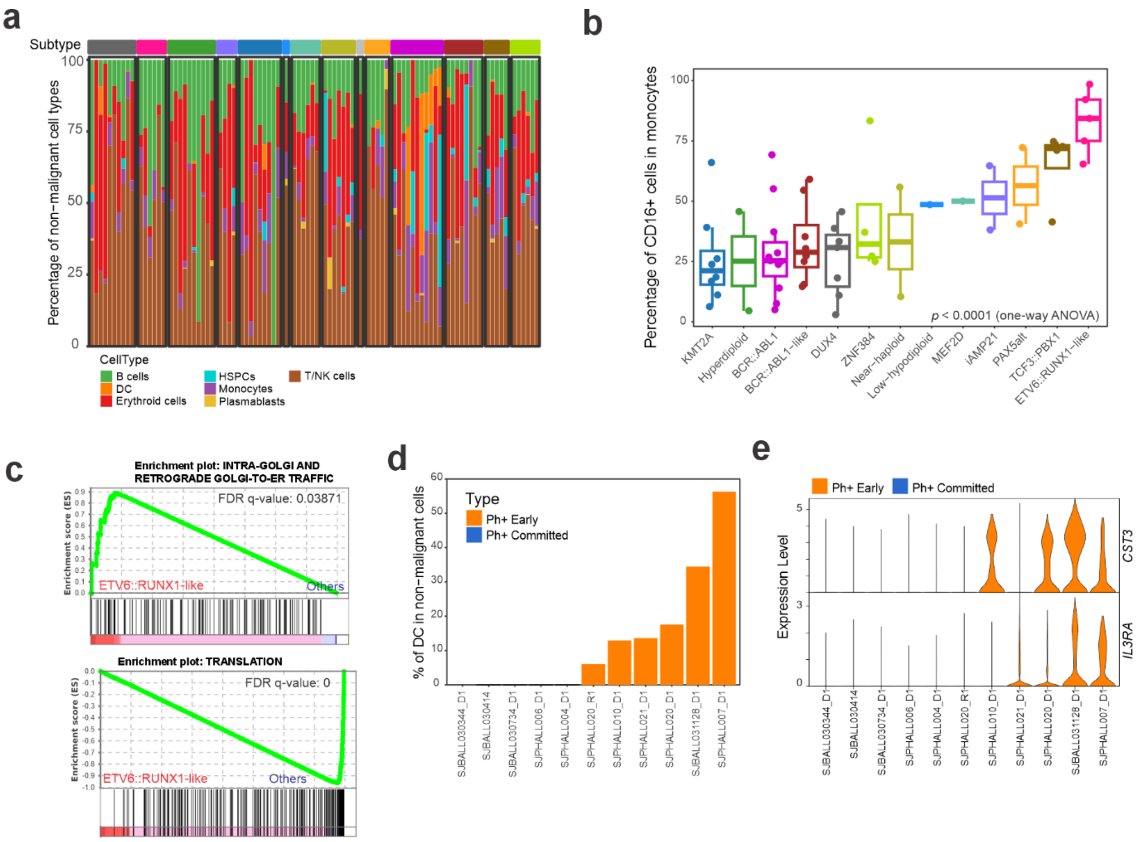

**Supplementary Figure. Single-cell analysis of non-malignant cells in B-ALL samples**. **a**) Schematic workflow of analysis of normal cells which include monocytes (Mono) and dendritic cells (DC), T cells and natural kill (NK) cells and normal B-cells. **b**) Percentage of CD16+ cells (non-classical monocytes) in all monocytes across different subtypes. *ETV6::RUNX1*-like samples had the highest amount (adjusted p value < 0.0001, one-way ANOVA). **c**) GSEA showing enrichment of ribosomal proteins in DC from *ETV6::RUNX1*-like samples. **d**) Percentage of DC in non-malignant cells in *BCR::ABL1* positive ALL patients. **e**) Violin plot showing DC-genes *BCR::ABL1* ALL samples.

### Supplementary References

1. Paulsson, K.*, et al.* The genomic landscape of high hyperdiploid childhood acute lymphoblastic leukemia. *Nat Genet* **47**, 672-676 (2015).
2. Woodward, E.L.*, et al.* Clonal origin and development of high hyperdiploidy in childhood acute lymphoblastic leukaemia. *Nat Commun* **14**, 1658 (2023).
3. Brady, S.W.*, et al.* The genomic landscape of pediatric acute lymphoblastic leukemia. *Nat Genet* **54**, 1376-1389 (2022).
4. Khabirova, E.*, et al.* Single-cell transcriptomics reveals a distinct developmental state of KMT2A-rearranged infant B-cell acute lymphoblastic leukemia. *Nat Med* **28**, 743-751 (2022).
5. O'Byrne, S.*, et al.* Discovery of a CD10-negative B-progenitor in human fetal life identifies unique ontogeny-related developmental programs. *Blood* **134**, 1059-1071 (2019).
6. Jackson, T.R., Ling, R.E. & Roy, A. The Origin of B-cells: Human Fetal B Cell Development and Implications for the Pathogenesis of Childhood Acute Lymphoblastic Leukemia. *Front Immunol* **12**, 637975 (2021).
7. Witkowski, M.T.*, et al.* Extensive Remodeling of the Immune Microenvironment in B Cell Acute Lymphoblastic Leukemia. *Cancer Cell* **37**, 867-882 e812 (2020).
8. Popescu, D.M.*, et al.* Decoding human fetal liver haematopoiesis. *Nature* **574**, 365-371 (2019).
9. Jardine, L.*, et al.* Blood and immune development in human fetal bone marrow and Down syndrome. *Nature* **598**, 327-331 (2021).
10. Roy, A.*, et al.* Transitions in lineage specification and gene regulatory networks in hematopoietic stem/progenitor cells over human development. *Cell Rep* **36**, 109698 (2021).
11. Oetjen, K.A.*, et al.* Human bone marrow assessment by single-cell RNA sequencing, mass cytometry, and flow cytometry. *JCI Insight* **3**(2018).
12. Ainciburu, M.*, et al.* Uncovering perturbations in human hematopoiesis associated with healthy aging and myeloid malignancies at single-cell resolution. *Elife* **12**(2023).
13. Setty, M.*, et al.* Characterization of cell fate probabilities in single-cell data with Palantir. *Nat Biotechnol* **37**, 451-460 (2019).
14. Granja, J.M.*, et al.* Single-cell multiomic analysis identifies regulatory programs in mixed-phenotype acute leukemia. *Nat Biotechnol* **37**, 1458-1465 (2019).
15. Mende, N.*, et al.* Unique molecular and functional features of extramedullary hematopoietic stem and progenitor cell reservoirs in humans. *Blood* **139**, 3387-3401 (2022).
16. Bueno, C.*, et al.* CD34+CD19-CD22+ B-cell progenitors may underlie phenotypic escape in patients treated with CD19-directed therapies. *Blood* **140**, 38-44 (2022).
17. Triana, S.*, et al.* Single-cell proteo-genomic reference maps of the hematopoietic system enable the purification and massive profiling of precisely defined cell states. *Nat Immunol* **22**, 1577-1589 (2021).
18. Kotliar, D.*, et al.* Identifying gene expression programs of cell-type identity and cellular activity with single-cell RNA-Seq. *Elife* **8**(2019).
19. Choudhary, S. & Satija, R. Comparison and evaluation of statistical error models for scRNA-seq. *Genome Biol* **23**, 27 (2022).
20. Korotkevich, G., Sukhov, V. & Sergushichev, A. Fast gene set enrichment analysis. *bioRxiv*, 060012 (2019).
21. Ben-Kiki, O., Bercovich, A., Lifshitz, A. & Tanay, A. Metacell-2: a divide-and-conquer metacell algorithm for scalable scRNA-seq analysis. *Genome Biol* **23**, 100 (2022).
22. Lambert, S.A.*, et al.* The Human Transcription Factors. *Cell* **172**, 650-665 (2018).

23. Aibar, S., *et al.* SCENIC: single-cell regulatory network inference and clustering. *Nat Methods* **14**, 1083-1086 (2017).
